# Cachd1 is a novel Frizzled- and LRP6-interacting protein required for neurons to acquire left-right asymmetric character

**DOI:** 10.1101/2022.05.16.492129

**Authors:** Gareth T. Powell, Ana Faro, Yuguang Zhao, Heather Stickney, Laura Novellasdemunt, Pedro Henriques, Gaia Gestri, Esther Redhouse White, Jingshan Ren, Weixian Lu, Rodrigo M. Young, Thomas A. Hawkins, Florencia Cavodeassi, Quenten Schwarz, Elena Dreosti, David W. Raible, Vivian S. W. Li, Gavin J. Wright, E. Yvonne Jones, Stephen W. Wilson

**Affiliations:** Cell and Developmental Biology, University College London, London, WC1E 6BT, UK; Wellcome Trust Sanger Institute, Wellcome Trust Genome Campus, Hinxton, Cambridgeshire, CB10 1SA, UK; Division of Structural Biology, Wellcome Centre for Human Genetics, University of Oxford, Roosevelt Drive, Oxford, OX3 7BN, UK; Department of Biological Structure, Health Sciences H-501, University of Washington, 1959 NE Pacific St., Seattle, WA 98195-7420, USA; The Francis Crick Institute, 1 Midland Road, London, NW1 1AT, UK; Institute of Ophthalmology, University College London, London EC1V 9EL, UK; St. George’s, University of London, Cranmer Terrace, SW17 0RE, London, UK; Department of Biology, Hull York Medical School, York Biomedical Research Institute, University of York, Wentworth Way, York, YO10 5DD, UK

**Author notes:** indicates joint authorship (see Author Contributions) (Gareth T. Powell, Ana Faro, Yuguang Zhao, Heather Stickney).

## Abstract

Neurons on left and right sides of the nervous system frequently show asymmetric properties but how these differences arise is poorly understood. Through a forward genetic screen in zebrafish, we find that loss of function of the transmembrane protein Cachd1 results in right-sided habenula neurons adopting left-sided character. Cachd1 is expressed in habenula neuron progenitors, functions symmetrically downstream of asymmetric environmental signals that determine laterality and influences timing of the normally left-right asymmetric patterns of neurogenesis. Unbiased screening for Cachd1 partners identified the Wnt co-receptor Frizzled7 and further biochemical and structural analysis revealed Cachd1 can bind simultaneously to Fzd proteins and Lrp6, bridging between these two Wnt co-receptors. Consistent with these structural studies, *lrp6* mutant zebrafish show symmetric habenulae with left-sided character and epistasis experiments with other Wnt pathway genes support an *in vivo* role for Cachd1 in modulating Wnt pathway activity in the brain. Together, these studies identify Cachd1 as a conserved novel Wnt-receptor interacting protein with roles in regulating neurogenesis and neuronal identity.

## Introduction

The nervous systems of bilaterian animals are frequently, perhaps universally, left-right (LR) asymmetric with respect to neuroanatomy, processing of information and control of behaviour (*1, 2*). Within vertebrates, the epithalamus shows evolutionarily conserved LR asymmetries (*3, 4*). In zebrafish, the epithalamic dorsal habenulae (dHb) comprise medial (dHbM) and lateral (dHbL) domains each containing distinct subtypes of projection neuron; the dHbL is larger on the left whereas the dHb_M_ is larger on the right (*5–7*). Functional asymmetry mirrors neuroanatomy; for instance, in young fish, light activates predominantly left-sided dHb_L_ neurons, whereas a higher proportion of right-sided dHb_M_ neurons respond to odour (*8*). Afferent innervation is also asymmetric with a subset of mitral cells innervating the right dHb and the photosensitive parapineal nucleus innervating the left dHb e. g. (*9–11*).

The development of epithalamic asymmetry is dependent upon sequential interactions between cell groups that coordinate lateralisation of circuit components (*12–14*). Although the developmental mechanisms by which asymmetry arises remain poorly understood, genetic analyses in zebrafish have revealed multiple roles for Wnt signalling during the establishment of lateralised circuitry. For instance, the habenulae of fish with compromised function of Axin1, a scaffolding protein in the ß-catenin degradation complex, have symmetric habenulae with right-sided character whereas the habenulae are symmetric with left-sided character in fish lacking function of the Tcf7l2 transcriptional effector of Wnt signalling (*15, 16*). Wnt signalling also impacts habenula size, as fish with compromised function of Wls, a protein involved in Wnt secretion, have small, but asymmetric habenulae (*17*). These and other studies (*18*) indicate that Wnt signalling regulates several distinct aspects of epithalamic development suggesting complex regulation of Wnt pathway activity during this process.

The Wnt signalling pathway is involved in a wide array of biological processes during embryonic development, throughout life and in many disease states, including cancer (*19*). Wnt ligands can activate several signalling cascades and pathway activation is tightly regulated at many different steps from ligand production to ligand/receptor interactions, cytoplasmic signal transduction and transcriptional activation/repression (*20–22*). Through studying the role of Wnt signalling in the establishment of brain asymmetry, here we identify Cachd1 as a novel transmembrane component of this highly conserved and multi-functional signalling pathway.

## Results

### *rorschach^u761^* mutants show symmetric habenulae due to a lesion in the *cachd1* gene

To identify novel genes involved in the establishment of brain asymmetry, we screened zebrafish embryos carrying ENU-induced mutations that alter asymmetric habenular expression of the *kctd12.1* gene (*16*) and identified the *rorschach^u761^* mutant (*rch*). In 4 dpf larvae homozygous for the *u761* mutation, *kctd12.1* expression (*5*) in the right habenula was increased to the level within the left habenula suggesting that both habenulae exhibit left-sided character (Fig. 1A). Other than this fully penetrant habenular phenotype, homozygous *rch* mutants were morphologically indistinguishable from wild types with normal asymmetry and laterality of the viscera.

**Fig. 1.**
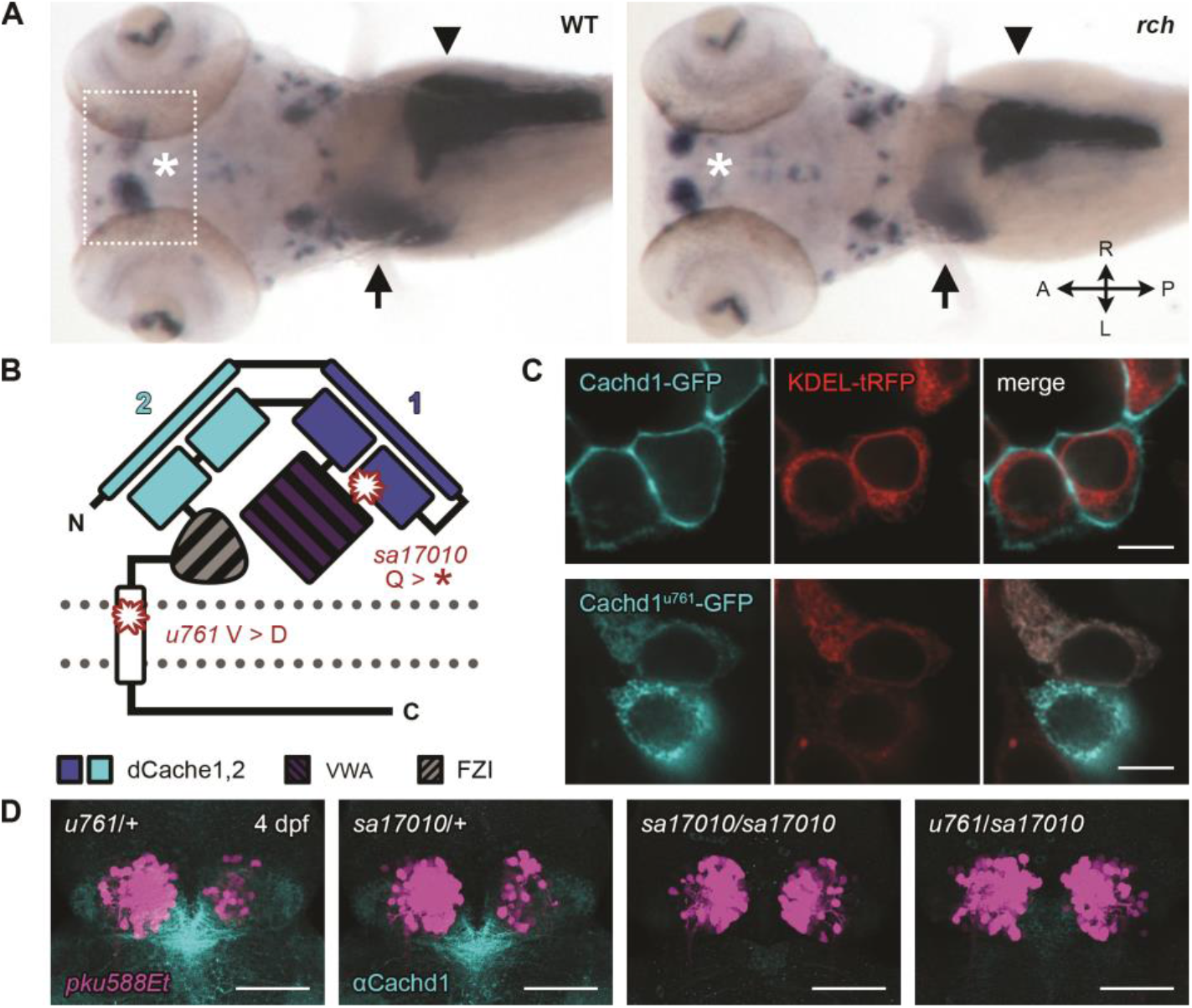
*cachd1* mutants show bilaterally symmetrical, ‘double left’ habenulae. (A) Dorsal views of wholemount *in situ* hybridisation 5 dpf wildtype sibling and *rorschach* (*rch/u761*) mutant larvae, stained with an asymmetric marker of the habenulae (*kctd12.1,* asterisk, box indicates approximate epithalamic region in D) and markers for liver (*selenop2,* arrow), pancreas (*prss1,* arrowhead), and ventral retina (*aldh1a3*). (B) Schematic of Cachd1 protein: two dCache domains (cyan and dark blue), a VWA domain (purple stripes), a Fzd binding domain (FZI, grey stripes), a transmembrane domain (white) and an unstructured cytoplasmic tail. Residues affected in *sa17010* and *u761* alleles are marked in red at approximate positions in primary sequence. (C) Fluorescence images of transfected HEK293T cells expressing constructs encoding GFP-tagged wildtype (top row, cyan) or *rch*/*u761* mutant Cachd1 (bottom row, cyan) and KDEL-tRFP (red) to mark the endoplasmic reticulum. Scale bar = 10 μm. (D) Dorsal views of brains of dissected 4 dpf transgenic siblings from a complementation cross with *sa17010* and *u761* alleles, stained with anti-Cachd1 antibody (cyan). *Et(gata2a:eGFP)pku588 (pku588Et)* expresses GFP in dHb_L_ neurons (magenta). Maximum projections of confocal z-stacks, scale bar = 50 μm.

Mapping using bulked segregant analysis followed by high resolution simple sequence length and single nucleotide polymorphism analyses placed the *rch* mutation in a 0.28 Mb interval on linkage group 6, encompassing *ak4, jak1, raver2* and *cachd1.* Sequencing identified a non-synonymous single base pair change in *cachd1* switching a nonpolar valine (V; GTT) at amino acid position 1122 to an acidic aspartic acid (D; GAT). *cachd1* encodes a 1290 amino acid type I transmembrane protein with dCache and von-Willebrand factor (VWA) domains; the V1122D missense mutation occurs within the transmembrane domain (Fig. 1B). To determine whether the V1122D substitution affects protein localization, we fused eGFP to the C-terminal end of wildtype and mutant Cachd1 proteins and examined localization of the fusion proteins in transfected HEK293 cells. As expected, wildtype Cachd1-eGFP fusion protein localized to cell membranes whereas the Cachd1^u761^-eGFP fusion protein remained cytoplasmic, co-localizing with the endoplasmic reticulum (Fig. 1C, see also Fig. S1A, B).

To confirm that the *u761* mutation in *cachd1* is causative of the symmetric habenular phenotype, we analysed phenotypes after abrogation of *cachd1* in other contexts. Embryos homozygous for a likely null mutation in *cachd1* (*sa17010*), that makes no detectable Cachd1 protein (by immunohistochemistry; Fig. S1C, Table S1), showed the same habenular double left-phenotype, as did transheterozygote *cachd1^u761^/cachd1^sa17010^* mutants (n = 10/10, Fig. 1D; n = 6/6, Fig. S2) and embryos injected with splice-blocking *cachd1* morpholinos (*kctd12.1,* n = 253/263, *kctd12.2*, n = 16/17, Fig. S3). Finally, habenular asymmetry was partially restored in homozygous *cachd1^u761^* mutants expressing exogenous, wildtype Cachd1 from a heat shock promoter prior to and during the period of habenular neurogenesis (*Tg*(*hse:cachd1,gfp*)*w160*, Fig. S4). Together, these results confirm that loss of Cachd1 function underlies the symmetric habenular phenotype.

### Cachd1 is expressed in neuroepithelial cells along the dorsal midline of the brain

To determine where and when *cachd1* is expressed within the brain, we performed fluorescent *in situ* hybridisation and immunohistochemistry using an antibody raised and purified against the extracellular domain of zebrafish Cachd1 (Fig. 2, Fig. S1). Prior to neuronal differentiation, *cachd1* is present broadly within the dorsal diencephalon (Fig. 2A), and co-localises with *dbx1b,* an early marker of habenula neuron precursors (Fig. 2A, B) (*23*). During the period of habenular neurogenesis (*24*), *cachd1*/Cachd1 expression becomes restricted to a proliferative neuroepithelial domain close to the midline adjacent to mature habenula neurons (Fig. 2C-E’, Fig. S5). Although the *cachd1* mutant only shows an overt mutant phenotype on the right side of the brain, we detected no obvious asymmetry in *cachd1*/Cachd1 expression until much later in development, long after habenula asymmetry has been established (Fig. S6). Early Nodal signalling-dependent brain (*25, 26*) and visceral (*27*) asymmetries were unperturbed in *cachd1* mutant embryos (Fig. S7). Taken together, these results suggest that *cachd1* functions locally within the progenitor domain that gives rise to habenula neurons.

**Fig. 2.**
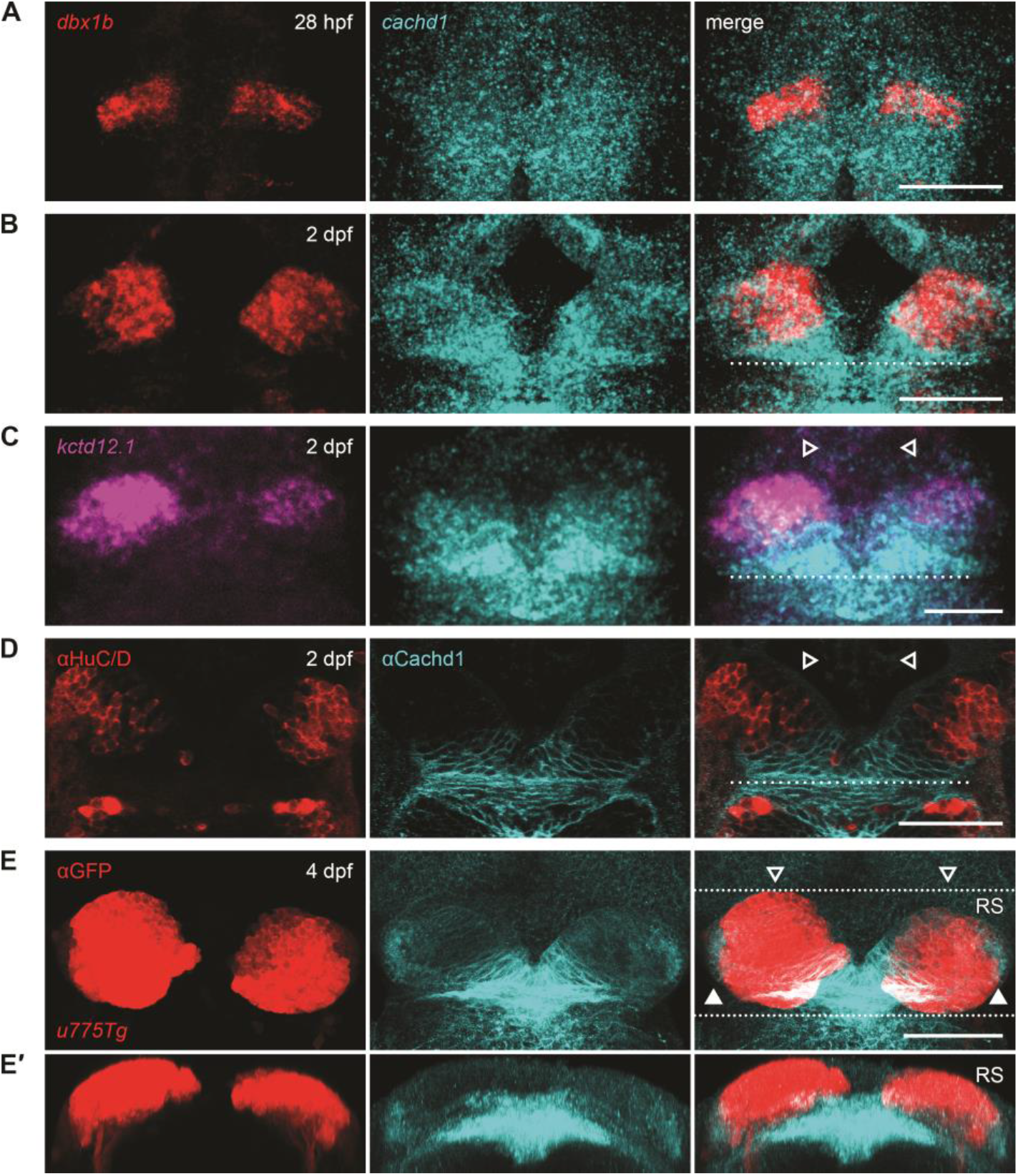
*cachd1* is expressed by presumptive habenula neuron progenitors. (A-C) Dorsal views of habenulae after double fluorescent *in situ* hybridisation with *cachd1* antisense riboprobe (cyan) and riboprobes for either the habenula neuron progenitor marker *dbx1b* (red) at 28 hpf (A) and 2 dpf (B), or the dHb_L_ marker *kctd12.1* (magenta) at 2 dpf (C). (D-E) Dorsal views of habenulae after immunohistochemistry with anti-Cachd1 antibody (cyan) co-stained with either anti-HuC/D antibody to mark differentiated neurons (red) at 2 dpf (D) or anti-GFP antibody (red) to mark habenula neurons expressing GFP in *Tg(110316_GFP)u775 (u775Tg)* embryos at 3 dpf (E). All images are maximum projections of confocal stacks, except for a single confocal slice in (D). A single dotted line in (B-D) indicates the approximate position of the posterior commissure; open arrowheads indicate the dorsal habenulae (C-E); closed arrowheads indicate the ventral habenulae (E, weakly expressing GFP). The dotted lines in (E) indicate the region (RS) represented in the transverse projection in (E’). Scale bars = 50 μm (A-E)

### Cachd1 functions in both habenulae to promote right-sided and/or suppress left-sided character

The right dorsal habenula (dHb) differs from the left dorsal habenula with respect to gene expression, organisation of synaptic neuropil and targeting of projection neuron connections (*5–7, 11, 28*); asymmetries of all these features in *cachd1* mutants were reduced or absent with the right habenula closely resembling the left habenula (Fig. 3A-B’ and Fig. S8). The dHb contain two broad sub-types of projection neuron present in different frequencies on right and left (*6, 7, 24, 28*). In the left dHb, neurons that project to the dorsal interpeduncular nucleus (dIPN; termed dHb_L_ neurons) are predominant whereas on the right, the majority of neurons project to the ventral IPN (vIPN; termed dHb_M_ neurons). Unlike in wildtypes, in *cachd1^u761^* mutants, DiI/DiD labelling revealed that the right dHb extensively innervated the dIPN, consistent with a higher proportion of right-sided dHb neurons adopting dHb_L_ character (n = 3 wildtype siblings, 8 *u761* mutants; Fig. 3A-B’). Together these results show that on the right side of the brain, Cachd1 function promotes dHbM and/or suppresses dHb_L_ character. However, they do not reveal whether Cachd1 has any function in determining the molecular character of the left side of the brain.

**Fig. 3.**
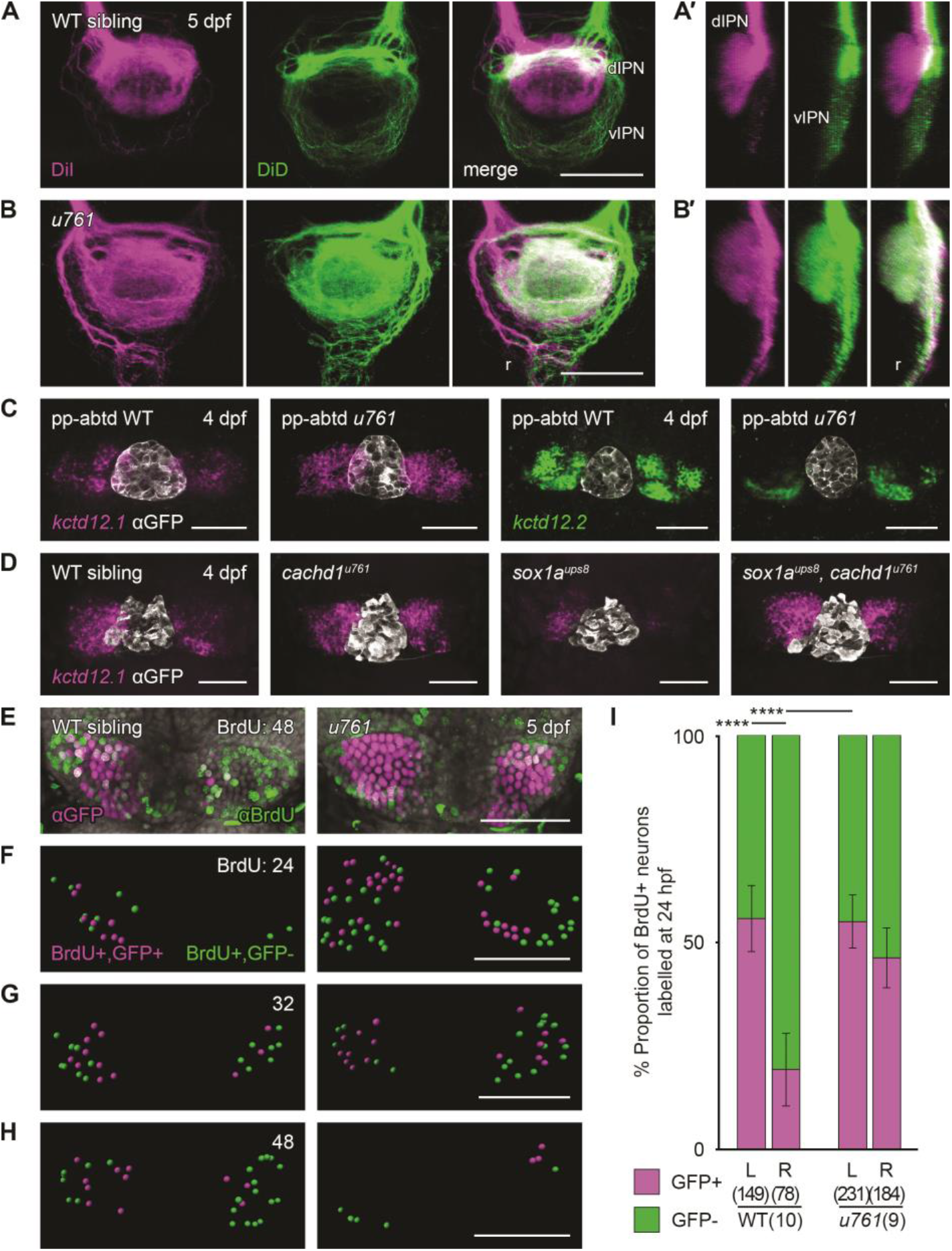
Loss of function of *cachd1* disrupts habenular efferent connectivity, is epistatic to removal of the parapineal signal and causes precocious neurogenesis. (A-B) Dorsal views and sagittal projections (A’-B’, dorsal left) of the IPN showing DiI (magenta) and DiD (green) labelling of left- and right-sided habenula neuron axon terminals predominantly innervating the dIPN and vIPN respectively, and raphe (r), in 5 dpf wildtype (A, n = 3) or *cachd1^u761^* mutant (B, n = 8) larvae. (C) Dorsal views of 4 dpf wildtype or *cachd1^u761^* mutant epithalami in which the parapineal was ablated prior to leftward migration (pineal complex marked by *zf104Tg, u711Tg* alleles with anti-GFP, white) after double FISH with *kctd12.1* (magenta; n = 26/29 wildtype siblings, 11/12 *cachd1^u761^* mutants) or *kctd12.2* (green; n = 19/23 wildtype siblings, 5/5 *cachd1^u761^* mutants). (D) Dorsal views of 4 dpf larvae from an epistasis cross of *cachd1^u761^* and *sox1a^ups8^* alleles after FISH with *kctd12.1* (magenta; pineal complex as C, white). n = 4 wildtypes, 3 *cachd1^u761^* mutants, 4 *sox1a^ups8^* mutants, 3 *sox1a^ups8^, cachd1^u761^* double mutants. (E) Dorsal views of *Et(gata2a:eGFP)pku588* wildtype or *cachd1^u761^* mutant habenulae incubated at 48 hpf with a pulse of BrdU to label newly born neurons, then processed for immunohistochemistry at 5 dpf with anti-GFP (magenta) and anti-BrdU (green) antibodies. DAPI counterstain marking nuclei (grey). (F-H) Segmentation of confocal stacks from *Et(gata2a:eGFP)pku588* wildtype or *cachd1^u761^* mutant larvae incubated at 24 (F), 32 (G) and 48 hpf (H) with a pulse of BrdU then processed at 5 dpf as in (E). Double positive cells are represented in magenta; BrdU-positive only cells are represented in green. Time of pulse indicated in top right corner. (I) Quantification of the proportion of BrdU-positive neurons that also expressed *Et(gata2a:eGFP)pku588* (magenta) in 5 dpf wildtype or *cachd1^u761^* larvae incubated with a pulse of BrdU at 24 hpf (all timepoints presented in Fig. S9). Error bars represent 95% confidence intervals. Total number of cells and larvae for each genotype indicated in axis label in brackets. Q’ test of equality of proportions (all timepoints, degrees of freedom = 15, *χ^2^* = 747.49, p = 1.39 × 10^-149^), *post hoc* pairwise comparisons using a modified Marascuilo procedure with Benjamini-Hochberg correction for multiple testing, **** p < 0.005. Scale bars = 50 μm (A-H)

The parapineal nucleus, a small group of cells present on the left side of the brain, is critical for the elaboration of most aspects of left-sided habenula character (*5, 7, 11, 29*). Consequently, if the parapineal is ablated (Fig. 3C) or fails to signal (Fig. 3D, *sox1a^ups8^* mutant) (*29*), the left dHb develops with right-sided character. To examine if the left-sided character of the habenulae in *cachd1* mutants is dependent upon parapineal signalling, we compared the effects of parapineal ablation between wildtype and *cachd1^u761^* mutant embryos. As expected, ablation of the parapineal in wildtype siblings led to reduced expression *kctd12.1,* which is normally expressed at high levels on the left (n = 26/29; Fig. 3C), and increased expression of *kctd12.2,* normally expressed at low levels on the left (n = 19/23; Fig. 3C). By contrast, the double-left habenular phenotype of *cachd1* mutants was unaffected by parapineal ablation (*kctd12.1:* n = 11/12, *kctd12.2:* n = 5/5; Fig. 3C). Similarly, in *cachd1^u761^, sox1a^ups8^* double mutants (n = 3/3; Fig. 3D) the *cachd1* mutant phenotype was epistatic to the *sox1a* mutant phenotype. Taken together, these results imply that Cachd1 functions on both sides of the brain to suppress left-sided character and/or promote right-sided character. As a corollary to this, it also implies that the role of the parapineal is to abrogate the function of Cachd1 within the left habenula.

Both the timing of neurogenesis and the environment into which habenula neurons are born influence their sub-type identity (*16, 24*). dHb_L_ neurons tend to be generated earlier than dHb_M_ neurons and neurogenesis is initiated earlier in the left habenula than in the right habenula (*24*). Furthermore, early born neurons differentiating on the left have a higher probability of adopting dHb_L_ character than neurons born on the right (*16*). To elucidate how Cachd1 impacts these asymmetries in neurogenesis, we performed birth dating experiments to assess both the extent of habenular neurogenesis at different stages and the timing of birth of dHb_L_ neurons, marked with the transgene *Et(gata2a:eGFP)pku588 (pku588Et,* Fig. 3E-I). These analyses showed that neurogenesis began early in *cachd1^u761^* mutants compared to wildtype, was symmetric on the left and right sides of the brain (Fig. 3F-G, Fig. S9) and diminished over time (Fig. 3H, Fig. S9). In addition, early born neurons in the right habenula of *cachd1* mutants had a higher likelihood of taking on dHb_L_-character than in wildtype embryos, as assessed by subsequent expression of the *pku588Et* transgene (Fig. 3I; Fig. S9) and the dHb_L_ marker *kctd12.1* (Fig. S10).

### CACHD1 binds to Wnt pathway receptors

Given that the phenotype of embryos homozygous for the *u761* allele is a consequence of the absence of Cachd1 on the cell surface, we reasoned that the extracellular domain of Cachd1 is likely essential for its function and performed experiments to identify cell surface proteins that could physically interact with this receptor. In cultured cells, CACHD1, which has homology to the α2δ family of auxiliary subunits of voltage-gated Ca^2+^ channels (VGCCs), can alter VGCC activity and compete with other α2δ proteins (*30, 31*). However, to date there is no evidence of the necessity of this interaction during development *in vivo* and so we took an unbiased screening approach to find other interacting partners.

We initially identified FZD7 as a potential binding partner in a Retrogenix Cell Microarray Technology screen using a human CACHD1 ectodomain (ECD) multimer as a prey protein (see Methods; Fig. S11). To validate the interaction, we tested binding of the FLAG-tagged CACHD1 prey to live, intact HEK293E cells expressing full length, eGFP-tagged FZD7 (FZD7-eGFP) by flow cytometry. We observed a strong shift of secondary antibody anti-FLAG phycoerythrin-conjugate (PE) fluorescence in eGFP-positive cells tested with CACHD1 prey, but not an unrelated prey protein (FLAG-tagged rat CD200R ectodomain), suggesting specific binding of CACHD1 (Fig. 4A, B, Fig. S11). We found that binding was significantly reduced by pre-incubation of FZD7-eGFP-transfected cells with OMP-18R5, an anti-human FZD7 monoclonal antibody (*32*) (Fig. 4A, Fig. S12; n = 3, one-tailed paired *t*-test, degrees of freedom = 2, *t* = 9.53, p = 0.0054). OMP-18R5 binds an epitope in the extracellular cysteine rich domain (CRD) of several related FZD receptors (*32*) and so inhibition of binding by OMP-18R5 suggests that the N-terminal domain of FZD7 contains the binding site for CACHD1.

**Fig. 4.**
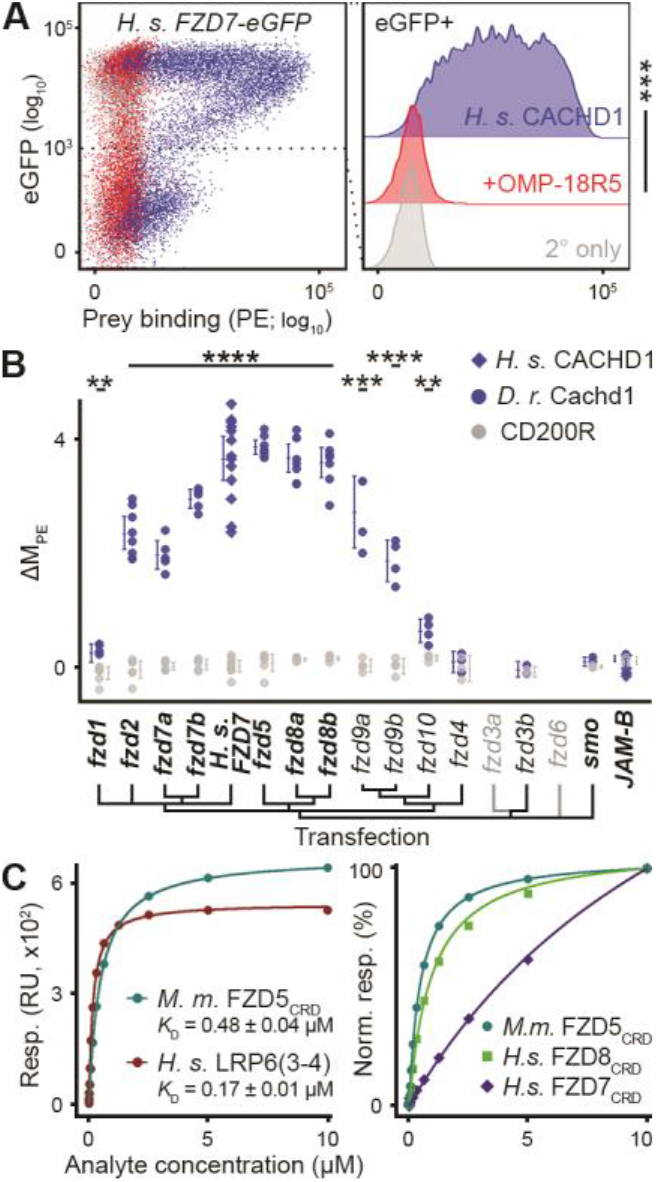
CACHD1 physically interacts with Wnt co-receptors LRP6 and FZD family members. (A, left) Representative scatter plot of flow cytometry testing binding of FLAG-tagged CACHD1 prey protein to human *FZD7-eGFP* transiently transfected HEK293E cells (detected by phycoerythrin (PE)-conjugated secondary antibody; A, right) without (blue) or with (red) pre-incubation with anti-Frizzled antibody OMP-18R5; secondary only negative control (grey). n = 3; one-tailed paired *t*-test (D. F. = 2, *t* = 9.53, *** p = 0.0054). (B) Dot plot of human (blue diamonds) or zebrafish (blue circles) CACHD1, or negative control CD200R (grey) prey protein binding (ΔM_PE_) to cells transiently transfected with eGFP fusion protein constructs indicated (transfections verified by antibody labelling in bold). Each dot represents a single experiment; horizontal bars denote the mean and error bars represent 95% confidence intervals. One way Welch test of means (Cachd1 prey v. CD200R prey, not assuming equal variances; F = 132.32, D. F_num_ = 30.00, D. F_denom_ = 34.67, p = 5.09 × 10^-28^), *post hoc* pairwise *t*-tests with non-pooled standard deviations, Benjamini & Hochberg correction for multiple testing; only significant differences between Cachd1 and CD200R prey for individual transfections are presented here for clarity, ** 0.05 > p > 0.01, *** 0.01 > p > 0.005, **** p < 0.005. (C) SPR-based determination of *K*_D_ for mouse CACHD1_ECD_ analyte binding to immobilised mouse FZD5_CRD_, human LRP6_P3E3P4E4_ (3-4, left panel), and normalised response curves for different CACHD1_ECD_:FZD_CRD_ interactions. RU, response units.

The most prominent feature of the N-terminal extracellular domain of the Frizzled family of receptors is the cysteine-rich domain (CRD). Because there is strong similarity of the CRD domains between Fzd proteins, we tested most zebrafish Frizzled family members for binding to Cachd1 prey using flow cytometry (quantified by comparing median PE fluorescence intensity of CACHD1 binding to eGFP-positive cells, 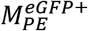, or untransfected eGFP-negative cells, 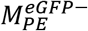, see Methods; summarised in Fig. 4B; Fig. S11). As expected, Cachd1 prey bound to cells transfected with eGFP fusion constructs of both zebrafish FZD7 orthologues: Fzd7a and Fzd7b, but was also able to bind to cells transfected with eGFP fusion constructs for almost all other members of the Frizzled family tested, except for Fzd4, Fzd3b and the control proteins. Interactions with Fzd1, Fzd2, Fzd7a and Fzd7b were also effectively inhibited by OMP18-R5 pre-incubation (Fig. S12). Furthermore, human CACHD1 prey protein was able to bind zebrafish orthologues of the Frizzled family and *vice versa* (Fig. S11C, D) suggesting strong conservation of the interactions.

To further characterise the interaction between CACHD1 and FZD proteins, we used surface plasmon resonance (SPR) to measure the binding affinity between purified recombinant mammalian CACHD1 and FZD orthologues. As expected, purified mouse Cachd1 extracellular domain analyte (CACHD1_ECD_) interacted with immobilised human FZD7_CRD_, albeit with low affinity (*K*_D_ ± 95% C.I.: 14.17 ± 2.18 μM, Fig. 4C, Fig. S13A, B). We also found that CACHD1_ECD_ interacts with mouse FZD5_CRD_ and human FZD8_CRD_ with much higher affinity (*K*_D_ ± 95% C.I.: 0.48 ± 0.04 μM and 0.95 ± 0.06 μM respectively, Fig. 4C, Fig. S13C, D, E).

Wnt ligands use FZDs and LRP5/6 coreceptors to initiate Wnt signalling (*20*). To test if CACHD1 could also interact with LRP6, we used immobilised human, membrane distal (LRP6_P1E1P2E2_) and membrane proximal (LRP6_P3E3P4E4_) fragments in SPR. CACHD1_ECD_ interacted with both LRP6 fragments, but with differing affinities: high affinity to the LRP6_P3E3P4E4_ fragment (*K*_D_ ± 95% C.I.: 0.17 ± 0.01 μM, Fig. 4C, Fig. S13F) and low affinity to LRP6_P1E1P2E2_ (*K*_D_ ± 95% C.I.: 5.86 ± 0.62 μM, Fig. S13G, H).

Taken together, these results demonstrate conserved interactions between CACHD1 and the Wnt co-receptors LRP6 and Frizzled family proteins.

### Structural characterisation of Cachd1 complex with FZD5 and LRP6

Guided by our *in vitro* measurements, we attempted co-crystallisation of CACHD1_ECD_ with FZD5CRD and LRP6P3E3P4E4. Crystals appeared in a PEG 4000 condition (see Methods) and diffracted to 4.7Å resolution. The structure was determined by molecular replacement using crystal structures of the CACHD1_ECD_:FZD5_CRD_ complex, previously determined in our laboratory (data not shown), and LRP6_P3E3P4E4_ (*33*) (PDB: 4A0P). The arrangement of the ternary complex in the crystal conforms to a C2_1_ space group with three complexes in an asymmetric unit (ASU). Refinement yielded complete structures of equivalent quality for all three copies (Table S2), of which one representative complex is depicted in Fig. 5A. CACHD1_ECD_ shows overall structural similarity to the α2δ1 auxiliary subunits of the voltage-gated Ca^2+^ channel Cav1.1 (*34*) (PDB: 5GJV, 778 Cα aligned at root mean square deviation = 4.4 Å), which contain two dCache domains and a Von Willebrand factor type A (VWA) domain. However, the Cachd1 structure reveals a novel addition to the C-terminal region of the ECD, which does not show any homology to known structures in Protein Data Bank by Dali search (*35*). This region interfaces with FZD5CRD (Fig. 5A) and we therefore term it the FZD interaction (FZI) domain. The two *α* helices of the N-terminal dCache domain (C-1) interact with the LRP6_P3_ propeller (Fig. 5A). Thus, CACHD1 serves as a crosslinking component in the ternary complex, independently binding to FZD5CRD and LRP6P3E3P4E4.

**Fig. 5.**
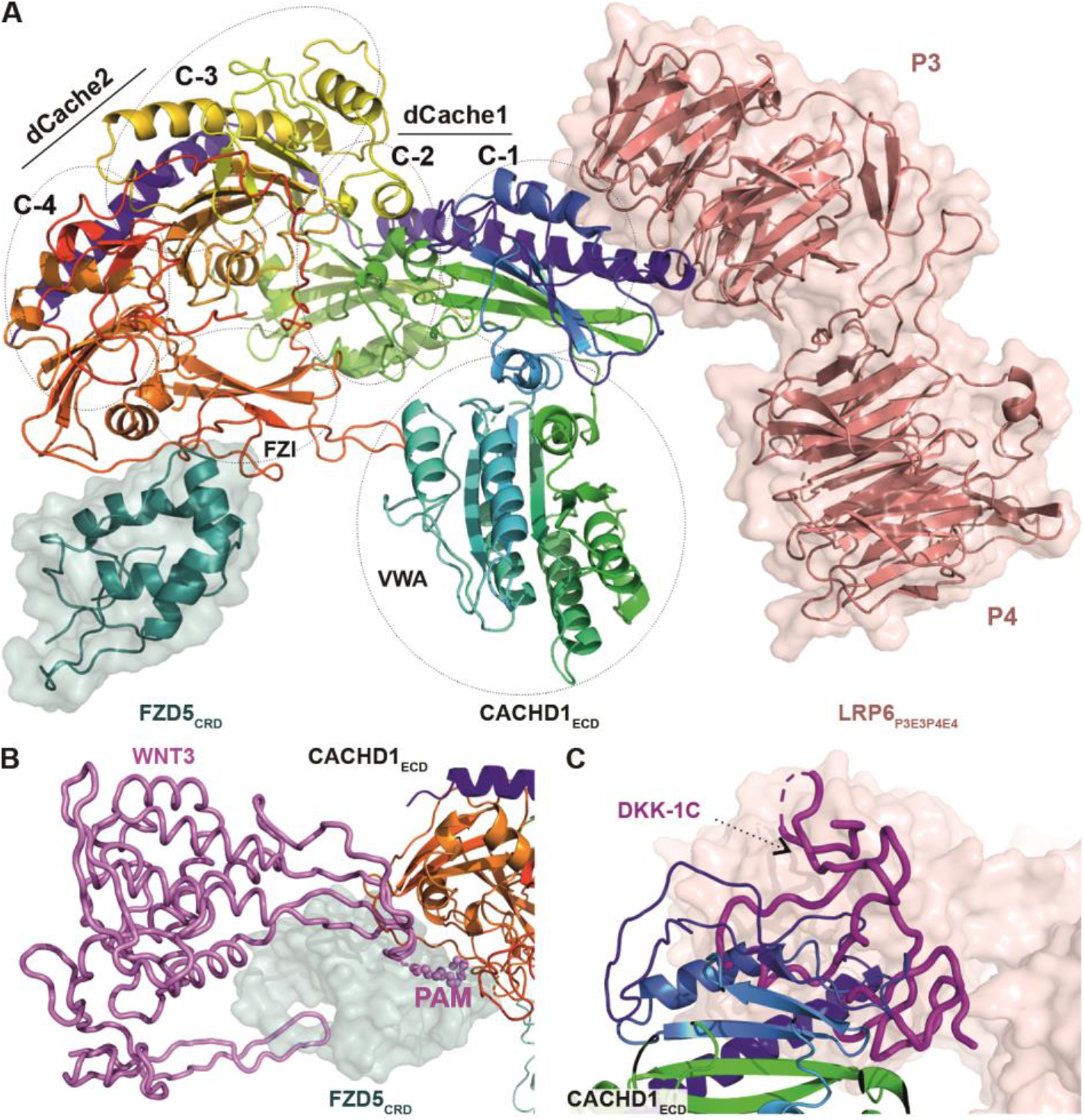
CACHD1 forms a ternary complex with FZD5 and LRP6. (A) Cartoon representation of mouse CACHD1_ECD_, (rainbow-colored from N-(blue) to C-(red) terminus) in complex with mouse FZD5_CRD_ (cartoon and surface in teal) and human LRP6_P3E3P4E4_ (cartoon and surface in salmon). The position of the four cache (C-1,2,3,4), VWA and FZD interaction (FZI) domains of CACHD1_ECD_ are indicated. (B) Superimposed structures of the FZD8:Wnt3 complex (PDB: 6AHY) with the FZD5_CRD_:CACHD1_ECD_ complex. Wnt3 is shown as a violet cartoon tube with palmitoleic acid (PAM) as spheres. (C) Superimposed structures of the LRP6:DKK1-C complex (PDB: 5FWW) with the CACHD1_ECD_:LRP6_P3E3P4E4_ complex. DKK1-C is shown as a magenta cartoon tube.

Structural superpositions show that the Cachd1 binding site on FZDCRD overlaps with the “thumb” and palmitoleic acid (PAM) lipid binding site (*36, 37*) required for the receptor-ligand interaction with Wnt (Fig. 5B). Functional studies have indicated that LRP6P3E3P4E4 harbours the primary binding site for Wnt3a (*38*) and also for the C-terminal domain of DKK-1 (DKK-1C), an inhibitor that competes with Wnts for binding to LRP5/6 (*20*). Crystal structures of LRP6_P3E3P4E4_:DKK-1 complexes (PDB: 3S2K, 3S8V & 5FWW) detail the interaction of the DKK-1 C-terminal domain with LRP6_P3_ (*39–41*). Superposition of our LRP6_P3E3P4E4_:Cachd1_ECD_ structure with the LRP6_P3E3P4E4_:DKK-1C complex (PDB: 5FWW) shows a steric clash between the CACHD1 C-1 helices and DKK-1C (Fig. 5C). This suggests that Cachd1 may also compete with Wnt3a for binding to the LRP6P3 propeller. Taken together our biophysical and structural analyses showed that CACHD1 is a novel binder to both members of the FZD family of Wnt receptors and the LRP6 co-receptors.

### *cachd1* genetically interacts with Wnt pathway genes

If Cachd1 functions together with Fzd and Lrp proteins during habenular development, then we might expect that abrogation of Fzd and/or Lrp6 function would also result in habenular asymmetry phenotypes. The Fzd family is large and we expect much redundancy between family members (*20*) and so we focussed on analysis of Lrp6 function in habenular development. We generated several predicted *lrp6* null alleles using CRISPR/Cas9 and found that homozygous mutant larvae showed a fully penetrant, symmetric double-left habenular phenotype, with visceral asymmetry unperturbed (*lrp6* mutants, n = 80/84, Fig. 6A, Fig. S14, Table S3). Consequently, loss of *cachd1* and *lrp6* function both result in symmetric habenulae with left-sided character, consistent with these two genes functioning in the same signalling pathway in the brain.

**Fig. 6.**
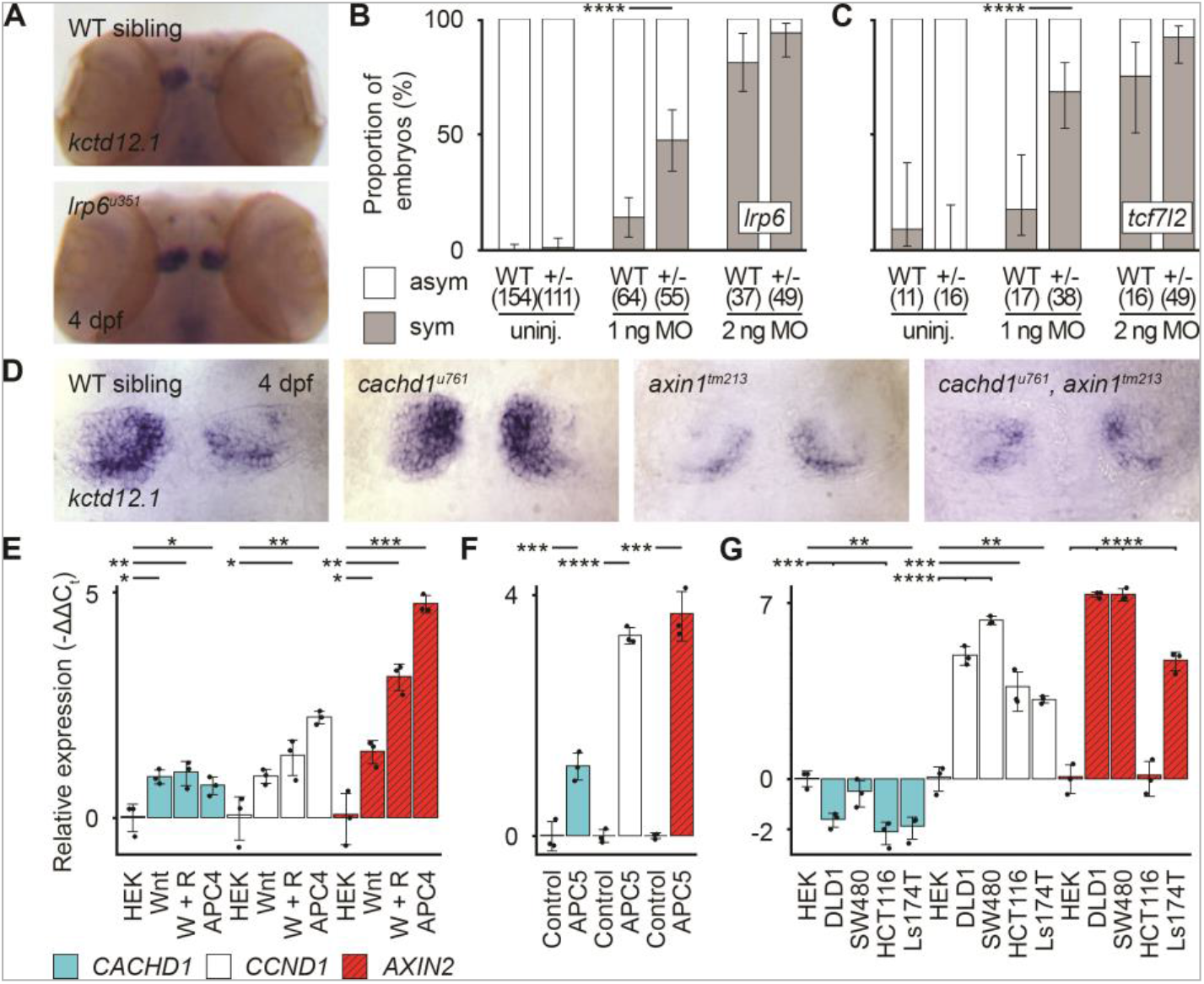
*cachd1* interacts genetically with Wnt pathway components. (A) Dorsal views of wholemount ISH 4 dpf wildtype sibling (n = 12) and *lrp6^u351^* mutant (n = 9) heads stained with *kctd12.1.* (B-C) Graphs showing the percentage of 4 dpf wildtype siblings and *lrp6^349/+^* larvae (B) or wildtype siblings and *tcf7l2^exI/+^* larvae (C) with (grey, sym) or without (white, asym) a symmetric bilateral left phenotype in uninjected larvae and larvae injected with a suboptimal or standard dose of *cachd1* morpholino (MO1). Error bars represent 95% confidence intervals of the proportion. Q’ test of equality of proportions (B: D. F. = 2, *χ^2^* = 18.71, p = 8.66 × 10^-5^; C: D. F. = 2, *χ^2^* = 7.93, p = 0.019) and *post hoc* modified Marascuilo procedure with Benjamini & Hochberg correction for multiple testing. **** represents p < 0.005. (D) Dorsal views of the habenulae of wholemount 4 dpf larvae from an incross of *cachd1^u761^* and *axin1^tm213^* mutants, showing expression of *kctd12.1.* n = 5 wildtypes, 6 *cachd1^u761^* mutants, 3 *axin1^tm213^* mutants and 3 *cachd1^u761^, axin1^tm213^* double mutants. (E-G) RT-qPCR data showing relative expression (-ΔΔC_t_) of *CACHD1* and known Wnt-responsive genes (*CCND1, AXIN2*) in (E) HEK293 (HEK) cells untreated, incubated with Wnt3a alone (Wnt) or Wnt3a and R-Spondin1 conditioned media (W + R), or stable *APC* mutant cells (APC4), (F) wildtype (Control) and APC mutant (APC5) mouse organoids and (G) colorectal cancer-derived cell lines. Data is presented as mean relative expression, compared to expression of *ACTB* (human) or *Hrpt1* (mouse) reference genes and untreated controls (HEK293 cells or wildtype organoid). Individual points represent biological replicates (each an average of three technical replicates), n = 3; error bars indicate 95% confidence intervals. One way Welch test of means (not assuming equal variances; A: F = 58.83, D. of F_num_ = 11.00, D. of F_denom_ = 9.41, p value = 3.03 × 10^-07^; B: F = 225.66, D. of F_num_ = 5.00, D. of F_denom_ = 5.16, p value = 5.12 × 10^-06^; C: F = 236.49, D. of F_num_ = 14.00, D. of F_denom_ = 11.33, p value = 7.67 × 10^-12^), *post hoc* pairwise *t*-tests with non-pooled standard deviations and Benjamini & Hochberg correction for multiple testing; only significant differences with control samples (HEK or Control) are presented here for clarity, * 0.1 > p > 0.05, ** 0.05 > p > 0.01, *** 0.01 > p > 0.005, **** p < 0.005.

We tested for a genetic interaction between *cachd1* and *lrp6* by injecting a validated *cachd1* morpholino into embryos from *lrp6^u349/+^* heterozygous outcrosses at a dose that results in a low frequency of symmetric habenular phenotypes in injected wildtype embryos. If the two genes interact genetically in the development of habenula asymmetry, then one might expect *lrp6* heterozygous embryos to be more sensitive to reduced Cachd1 function than their wildtype siblings. Indeed, we observed that heterozygous *lrp6^u349/+^* larvae were approximately three times more likely to be bilaterally symmetric for habenular *kctd12.1* expression than wildtype siblings when injected with a low dose (1 ng) of *cachd1* splice-blocking morpholino (MO1, Fig. 6B; bilateral phenotype proportion ± 95% confidence interval, n: WT: 0.14 ± 0.09, 64 larvae; *lrp6^u349/+^*: 0.47 ± 0.13, 55 larvae; Marascuilo procedure for comparing multiple proportions and Benjamini-Hochberg correction, degrees of freedom = 2, p = 0.0002). To confirm this difference was not due to morpholino efficacy, we injected a standard effective dose (2 ng) in parallel: a high degree of bilateral symmetry was observed in both genotypes (WT: 0.81 ± 0.13, 37 larvae; *lrp6^u349/+^*: 0.94, 95% CI: 0.68-0.94, 49 larvae) and only a single larva with a bilateral phenotype was observed in uninjected controls (WT: 0/154 larvae, *lrp6^u349/+^*: 1/111). These results are consistent with Cachd1 and Lrp6 functioning in the same developmental pathway during establishment of habenular left-right asymmetry.

Genetic interactions between *cachd1* and two other Wnt pathway genes implicated in habenular development (*tcf7l2* and *axin1*) were also examined (*15, 16*). Tcf7l2 is a transcriptional effector of Wnt signalling and loss of *tcf7l2* function results in symmetric habenulae with double-left character (*16*). *tcf7l2^exI/+^* heterozygotes show a wildtype habenular phenotype, however when Cachd1 levels were reduced in *tcf7l2^exI/+^* heterozygotes through injection of low dose *cachd1* morpholino, many larvae showed a symmetric, double-left phenotype (Fig. 6C; WT: 0.18, 95% CI: 0.06-0.41, 17 larvae; *tcf7l2^exI/+^*: 0.68, 95% CI: 0.53-0.81, 38 larvae; Marascuilo procedure for comparing multiple proportions and Benjamini-Hochberg correction, degrees of freedom = 2, p = 0.0001). This indicates that reduced activity of both genes results in a phenotype comparable to that seen when either gene alone is fully abrogated.

Axin1 is a scaffolding protein in the β-catenin degradation complex and compromised *axin1* function results in symmetric habenulae with double-right character (*15*), a phenotype opposite to that of *cachd1* mutants. When we generated *axin1^tm213^/cachd1^u761^* double mutants, they exhibited the *axin1* mutant phenotype (as assessed by expression of *kctd12.1,* Fig. 6D, *kctd8, scl18b* and *vachtb*, data not shown; n = 15/15); consequently, compromised Axin1 function is epistatic to loss of Cachd1 function. This is consistent with Axin1 functioning downstream of Cachd1 and the Fzd/Lrp6 receptor complex.

Wnt signalling often regulates expression of Wnt-pathway genes through positive and negative feedback mechanisms (*20*). Indeed, the spatially localised expression of *cachd1* along the roofplate and in the dorsal epithalamus is similar to that of other Wnt pathway genes such as the ligand encoding *wnt1, wnt3a* and *wnt10* genes and known Wnt pathway targets *axin2* and *lef1* (Fig. S15). We directly tested whether *CACHD1* is itself a target of Wnt signalling, using qPCR to assess *CACHD1* expression in cultured HEK293 cells treated with Wnt3a, or Wnt3a+RSpondin1 conditioned media, or a stable HEK293 cell line with a mutation in *APC* (named APC4) (*42*). *CACHD1* showed a similar level of transcriptional response to enhanced Wnt pathway activity as other Wnt target genes (Fig. 6E) suggesting that Cachd1 may be involved in Wnt signalling in other epithelial cell types. Similarly, *Cachd1* expression was upregulated in murine *Apc* mutant organoids (APC5, Fig. 6F). By contrast, cells derived from colorectal cancers (*APC* mutants: DLD1, SW480; β-catenin mutants: HCT116, Ls174T) showed downregulated *CACHD1* expression.

Together, these results provide compelling evidence to suggest that the structural interactions we have demonstrated are pertinent to Cachd1 function in the developing brain and that Cachd1 is a novel modulator of the Wnt signalling pathway, potentially functioning in many contexts.

## Discussion

Our results identify CACHD1 as a novel Wnt pathway component that bridges FZD and LRP6 Wnt co-receptors and functions in the developing brain and potentially other Wnt pathway contexts (*43*).

We have demonstrated evolutionary conserved interactions between CACHD1 and multiple FZD receptors through a previously unidentified FZI domain. Our crystal structure suggests that this domain could potentially compete with Wnts binding to FZDs through their PAM moiety. Similarly, CACHD1 binding to LRP6_P3E3P4E4_ through the first dCache domain suggests potential competition with either Wnt ligands or the Wnt inhibitor, DKK.

The simultaneous binding of Cachd1 to both Fzd and Lrp6 co-receptors suggests it may activate downstream signalling by clustering of the cytoplasmic signalling apparatus, as observed with artificial ligands (*44*). A role for Cachd1 in activating signalling would be consistent with the observed similarity of habenular phenotype in *cachd1, lrp6* and *tcf7l2* mutants, where the latter is thought to be due to a loss of pathway activation (*16*), and contrast to the phenotype of *axin1* mutants, in which the pathway is overactivated (*15*). However, the complexity of Wnt signalling frequently confounds simple interpretations and further studies are required to determine the *in vivo* signalling consequences of Cachd1/Fzd/Lrp6 interactions. For instance, the binding of Cachd1 to Fzd/Lrp receptors may simultaneously activate signalling and outcompete binding of Wnts, indirectly impacting activity of the pathway that might otherwise be set by the secreted ligand. Indeed, the relative differences in binding affinity to different Fzd proteins may render the consequences of Cachd1/Fzd/Lrp6 interactions highly context dependent.

Our study suggests that asymmetric Cachd1-dependent modulation of Wnt signalling leads to lateralisation of habenula neurons by altering both timing of neurogenesis and the probabilistic selection between alternate neuronal fates. We show that Cachd1 is present and can function on both sides of the brain but its activity on the left is antagonised by an unknown signal(s) from the parapineal. During habenular development, as in many other contexts, Wnt signalling appears to function at multiple stages and in multiple processes, from proliferation, through timing of neurogenesis, to acquisition and maintenance of neuronal identity (this study; (*15–18*)). It is largely unclear how this complexity of pathway activity and outcome is effected and an attractive possibility is that context-dependent activity of Cachd1 may contribute to this poorly understood aspect of Wnt signalling.

## Acknowledgements

We thank many colleagues for support and advice during the course of this project, staff at Diamond Light Source for assistance with X-ray data collection, Dr Austin Gurney for supplying the OMP-18R5 antibody, Jim Freeth, Mark Aspinall-O’Dea, Karen Williams and Natalia Guardiola at Charles River Discovery Research Services UK Limited for Cell Microarray technology and the UCL Fish facility for fish husbandry.

## Author Contributions

The senior authors wish to emphasise that all four lead authors made equally important contributions to this study and are happy for individuals to list the joint authors in whichever order they wish on CVs and other documents.

GTP devised and performed experiments (generated/gathered reagents, protein interactions/immunocytochemistry using flow cytometry, immunohistochemistry, *in situ* hybridisation chain reaction, *lrp6* mutagenesis, morpholino injections), analysed data, contributed to writing the paper, prepared figures. YZ devised and performed experiments (structural biology, protein interactions using surface plasmon resonance), analysed data, prepared figures. AF devised and performed experiments (*in situ* hybridisation, immunohistochemistry, BrdU pulse-chase, parapineal ablations, lipophilic dye labelling, epistasis, morpholino injections), analysed data. HS devised and performed experiments (ENU mutagenesis and screening, genetic mapping, gene identification, morpholino injections, *in situ* hybridisation, immunohistochemistry, lipophilic dye labelling, transgenesis and heat shock experiments, tissue culture and transfection), analysed data. LN devised and performed experiments (qRT-PCR, generated organoids), analysed data. PH devised and performed experiments (parapineal ablations and *in situ* hybridisation), analysed data. GG designed and performed experiments (phenotype characterisation). ER-W performed experiments (epistasis and *in situ* hybridisation). JR analysed data (structural phasing/model building). WL generated reagents for research (tissue culture/protein production). RMY, HS, TAH, FC and QS undertook the screen that isolated the *rorschach* mutant. ED generated reagents for research (transgenesis). DWR and VWSL provided funding for research (supported HS and LN respectively). GJW devised experiments and provided funding for research (supported GTP). EYJ provided funding for research (supported YZ, JR, WL), devised experiments, contributed to writing the paper. SWW provided funding for research (supported GTP, AF, HS, PH, ER-W, RMY, TAH, QS, FC, ED), devised experiments, wrote the paper.

## Data and materials availability

Further information and requests relating to zebrafish resources and reagents, including mutants generated in this study, should be directed to Steve W. Wilson, and those relating to structural biology and biochemistry, to E. Yvonne Jones.

## Materials and Methods

### Zebrafish husbandry and fish lines

Zebrafish were maintained in a designated facility according to UK Home Office and local regulations, on a 14h/10h light:dark cycle. Embryos were routinely stored in fish system water supplemented with methylene blue, or E3 embryo medium at 28°C. Zebrafish experiments and husbandry in the United States of America followed standard protocols in accordance with University of Washington Institutional Animal Care and Use Committee guidelines.

### Generation of mutant and transgenic lines

The *u761* mutant was generated by ENU mutagenesis. Mutations were induced in wild-type male AB/TL fish by four rounds of 3 mM ENU treatment as previously described (*45*).

The *sa17010* allele of *cachd1* was acquired from the Zebrafish Mutation Project (*46*).

An allelic series of predicted *lrp6* nonsense mutants (*u348, u349, u350* and *u351*) were recovered from founders mutated using CRISPR/Cas9. Briefly, *in vitro* transcribed, capped *cas9* mRNA and sgRNAs (prepared as described in (*47*) using T4 DNA polymerase, New England BioLabs, Ipswich, MA, USA and mMessage mMACHINE, Ambion, Austin, TX, USA) complementary to exon 2 of *lrp6* (sg1: GGCCAACGCCACGCTGGTGA, sg2: GGCCAGACCGGAGATGACGG; Table S4) were microinjected into the cell of 1 cell stage embryos. Injected fish were raised to adulthood, and genotyped for mosaicism of exon 2 using high resolution melting analysis (HRMA). Fish with a high degree of mosaicism were prioritised for outbreeding to generate F1s which were subsequently genotyped using headloop PCR combined with Sanger sequencing to identify alleles of interest (*48*) (Tables S3 and S5).

The *Tg(HSE:cachd1, GFP)w160* and *Tg*(*110316_GFP*)*u775* lines were generated by Tol2-mediated mutagenesis. Briefly, 1-cell zebrafish embryos were co-injected with *pTol2 HSE:cachd1*, *GFP* (*w160Tg*) or *pTol2 gng8:GFP* (*u775Tg*) construct (25-50 pg; see below) and capped *transposase* mRNA (40 pg) and the embryos raised to adulthood. Offspring of the injected, adult fish were screened for germline transmission of the transgene and their progeny raised.

### Cloning and genotyping of *u761*

Having used a combination of backgrounds to generate our F2s, we mapped *u761* in F3 embryos. We used bulked segregant analysis (*49*) followed by high resolution SSLP and SNP analyses to localize *u761* to a 0.28 MB interval on LG6 between a SNP in the first coding exon of *ak4* (2/5212 recombinants; *ak4* e1 primers; see Table S5) and an SSLP in intron 8-9 of *cachd1* (1/5212 recombinants; *cachd1* i8-9 primers; see Table S5). Sequencing of *cachd1* cDNA revealed a T to A transversion in the 24th exon of *cachd1* that causes a valine to aspartic acid amino acid substitution in its transmembrane domain (reverse strand 6:31607781 T>A, 1122V>D, Zv11 assembly). Mutants were subsequently genotyped with DCAPs primers (Table S5, *u761*-AloI primers) and the restriction enzyme AloI, which cuts the mutant allele, and then more routinely by KASP assay (see below).

### Morpholino knockdown

Two non-overlapping morpholino antisense oligonucleotides for *cachd1* (MO1: GTGTATTTTCCTACCTGCATGGTGA; MO2: AGGGATGATGTCTAACTCACCTGCT) were obtained from GeneTools (Philomath, OR, USA) and microinjected into the yolks of 1-cell stage zebrafish embryos in 1 nL volumes for a total dose between 4 ng and 0.5 ng, depending upon the experiment.

### DNA extraction, KASP and HRMA genotyping

Embryos or larvae were lysed at 95°C in 25 mM KOH, 0.2 mM EDTA for 30 minutes, cooled to 4°C and briefly vortexed to disrupt remaining tissue. The lysate was briefly spun in a centrifuge to collect, then neutralised with an equal volume of 40 mM Tris-HCl, pH 5.

KASP or HRMA genotyping of DNA lysates was performed as per manufacturer’s instructions, using either 2X KASP Master Mix with standard ROX (LGC Biosearch Technologies, Hoddesdon, UK) or 2X Precision Melt Supermix (Bio-Rad, Hercules, CA, USA), respectively, and a Bio-Rad CFX96 qPCR machine (see Table S5 for primer details).

Melting curves from HRMA genotyping were analysed using Precision Melt Analysis software (version 1.2; Bio-Rad).

### cDNA and plasmid constructs

Total RNA was extracted and purified from pools of 10-20 embryos (wildtype, *rch*, or *cachd1* MO-injected, depending on experiment) with TRIzol reagent (Life Technologies, Grand Island, NY, USA) according to the manufacturer’s instructions. cDNA was then produced using the SuperscriptIII first strand synthesis system for RT-PCR (Life Technologies).

*pCS2+ cachd1-eGFP and pCS2+ cachd1^u761^-eGFP:* Full-length *cachd1* and *cachd1^u761^* were amplified from cDNA with Phusion DNA polymerase (New England BioLabs) and primers tagged with SalI and SacII restriction enzymes sites (see Table S5 for primer sequences). The resulting PCR fragment and the *peGFP-N1-1* vector were sequentially digested with SalI and SacII prior to ligation with Quick ligase (New England Biolabs) to make a *pcachd1-eGFP-N1-1* plasmid. The *pcachd1-eGFP-N1-1* construct was then cut with SalI and HpaI and the fragment containing *cachd1*-*eGFP* cloned between the SalI and SnaBI sites of the *pCS2*+ vector.

*pTol2 HS:cachd1,GFP:* Full-length *cachd1* was amplified from cDNA using high-fidelity Phusion DNA polymerase (New England Biolabs) and then phosphorylated with PNK (New England BioLabs; see Table S5 for primers). The phosphorylated fragments were then cloned into the StuI site of a *pCS2*+ vector treated with Antarctic Phosphatase (New England Biolabs) to prevent recircularization. The resulting vectors were cut with BamHI and SnaBI and the *cachd1*-containing fragments cloned into the BamHI and EcoRV sites of the *pTol2 HSE:GFP* vector to obtain the *pTol2 HSE:cachd1,GFP* construct for injection.

*pTol2 gng8:GFP:* a 3060 bp promoter region of the *gng8* gene was amplified by PCR (see Table S5 for primer details) and cloned into a TOPO-TA vector. This fragment was subcloned into *pEGFP-N1,* upstream of the eGFP open reading frame, and the subsequent *gng8:eGFP* fragment cloned into *pTol2*.

For flow cytometry protein production, the coding sequence for the ectodomain of human and zebrafish *CACHD1* (truncated before the transmembrane domain at P1095/P1108 respectively) was codon optimised for HEK cells and synthesised by GeneArt (Thermo Fisher Scientific, Waltham, MA, USA). These fragments had NotI and AscI target sequences at the 5’ and 3’ ends, respectively, for subcloning into prey protein and ectodomain bait protein production vectors.

Human and zebrafish *CACHD1* prey protein expression constructs (ectodomain fused to a COMP domain, β-lactamase domain and FLAG tag) and zebrafish *cachd1* ectodomain production constructs (ectodomain fused to hexahistidine and BirA ligase peptide substrate tags) were prepared by NotI/AscI restriction enzyme double digest (New England Biolabs) of pTT3-based vector backbones (*50*) (Addgene IDs 71471 and 36153) and shuttle vectors containing the synthesised fragments, followed by ligation with T4 ligase (New England Biolabs). The resulting constructs were screened by Sanger sequencing to confirm correct in-frame insertion.

To create human and zebrafish *FZD-eGFP* bait protein constructs, IMAGE consortium clones (*51*) (see Table S6 for details) were used as templates in PCR reactions to generate full length inserts (including the seven transmembrane domains) with NotI and AscI target sequences at the 5’ and 3’ ends, respectively, except for *fzd4 and fzd9a* where the insert was synthesised by GenScript (Piscataway, NJ, USA) as no complete full length clone was available. *fzd1* and *fzd8b* both had NotI/AscI restriction sites in the respective coding sequences, so fusion PCR was used to generate full length inserts with synonymous mutations in the recognition sequences (see Table S5 for primer sequences). The PCR products were purified using a Qiaquick PCR purification kit (Qiagen, Hilden, Germany) and then digested with NotI/AscI (New England Biolabs) and ligated to a pTT3 vector containing eGFP (see below). The resulting constructs were verified by Sanger sequencing.

The pTT3-eGFP vector was constructed by replacing the C-terminal tag encoding region of a bait protein vector (*52*) (Addgene ID 36150) with eGFP. The bait protein vector was digested with AscI/BamHI (New England Biolabs) to remove the tag encoding region and then ligated to an eGFP insert generated by PCR using primers with AscI and BamHI tails (see Table S5 for primer sequences). The resulting vector was verified by Sanger sequencing to ensure in-frame insertion of the eGFP coding sequence.

All constructs used for producing proteins for surface plasmon resonance and crystallography were based on the mammalian stable expression vector pNeoSec (*53*). Mouse Cachd1 extracellular domain (UniProt: Q6PDJ1, residues D50–S1107) was derived from IMAGE clone 6834428 (Table S6; Source BioScience). Mouse Fzd5 cysteine rich domain (UniProt: Q9EQD0, residues A27-T157), human FZD7 CRD (UniProt: O75084, residues Q33-G170), human FZD8 CRD (UniProt: Q9H461, residues A28-T158) were synthesized (Genscript). Human LRP6 P1E1P2E2 (UniProt: O75581, residues A20-P630) and P3E3P4E4 (residues V629-G1244) domains were described previously (*44*).

### Tissue culture and cell transfection

HEK293T, APC4 (APC4 line was generated from HEK293T cells by CRISPR targeting APC with truncation at 1225 a.a.) (*42*), SW480, Ls174T, DLD1 and HCT116 were maintained in DMEM GlutaMAX (GIBCO) supplemented with 5% foetal bovine serum (FBS) (GIBCO), 100 U/mL penicillin (GIBCO) and 100 mg/mL streptomycin (GIBCO). All cells were maintained at 37°C in an incubator with 5% CO2. Cells were seeded in plates 24 hr before transfection, and plasmids were transfected using polyethylenimine (PEI; Polysciences, Warrington, PA, USA) or Fugene 6 (Promega) according to the manufacturer’s instructions.

HEK293T cells were simultaneously transfected with 1 μg pCS2Cachd1-eGFP or pCS2Cachd1_V1122D-eGFP and 1 μg KDEL-tRFP plasmid using Fugene 6 transfection reagent (Promega). eGFP/tRFP expression was confirmed 24 hours post transfection and the cells fixed at 42 hours post transfection and imaged on a spinning disk microscope (see below).

For protein production and flow cytometry experiments, suspension cultures of HEK293E or HEK293-6E cells were transfected using linear PEI:plasmid complexes, incubated for between 2-6 days and then harvested by centrifugation. The resulting cells or conditioned media were then used for downstream experiments (*50*). Briefly, HEK293E or HEK293-6E cells were maintained in suspension cultures in Freestyle 293 Expression Media (Gibco, Waltham, MA, USA) supplemented with heat-inactivated fetal calf serum (1%) and G418 (geneticin, 50 μg/mL; Sigma-Aldrich, St. Louis, MO, USA), routinely maintained at densities between 2.5 × 10^5^ and 4 × 10^6^ cells/mL in Erlenmeyer flasks (Corning, Corning, NY, USA) in a humidified orbital shaker at 37°C, 5% CO_2_. One day before transfection, cultures were split down to 2.5 × 10^5^ cells/mL in standard media or, in the case of biotinylated protein production, media supplemented with D-Biotin (100 μM). Plasmids for transfection were prepared using PureLink HiPure Plasmid Maxiprep kit, as per manufacturer’s instructions (Thermo Fisher Scientific), and resuspended in ddH_2_O at 1 mg/mL. For each transfection, purified plasmid was mixed with linear 25 kDa PEI (Polysciences) at a ratio of 1 μg DNA:2.2 μg PEI (per 5 × 10^6^ cells) in unsupplemented Freestyle 293 Expression Media (1/10^th^ culture volume), vortexed and left to stand for 5 minutes at room temperature to allow complexes to form, before mixing into the cell cultures (e. g. to transfect a 50 mL culture with density 5 × 10^5^ cells/mL, 50 μL of plasmid was mixed with 110 μL PEI 1 mg/mL in water, in 2 mL media). In the case of biotinylated protein production, cells were co-transfected with an additional secreted BirA ligase plasmid (*50*) (Addgene ID 64395) included in the transfection mixture at a ratio of 10 μg DNA:22 μg PEI: 1 μg BirA. Transfected cultures were incubated for approximately 2 (for flow cytometry) or 6 days (protein production) before harvesting by centrifugation (200 × g or 3200 × g, respectively) to separate cells from conditioned media.

Proteins used in surface plasmon resonance and crystallography were derived from stable cell lines established by G418 selection (1 mg/mL, Sigma) of transfected HEK293S GnTI(-) cells (*54*).

### Protein production and purification

Conditioned media was harvested from transfected cultures, pooled and filtered through 0.2 μm filters and stored at 4°C until use.

Prey protein transfections were quantified by β-lactamase assay, measuring the turnover of nitrocefin substrate by changing absorbance at 485 nm over time (*50*), then normalised by dilution.

Biotinylated bait ectodomain transfections were dialysed against PBS using SnakeSkin dialysis tubing (molecular weight cut-off 10,000 Da; Thermo Scientific) and several buffer changes (approximately 25 – 30 L in total). Biotinylated protein concentration was quantified by ELISA, using streptavidin-coated microplates and a monoclonal antibody to detect the CD4d3+4 tag (*50*) (Nunc Immobilizer, Thermo Fisher Scientific).

Unbiotinylated ectodomain transfections were collected and quantified by ELISA using nickel-coated microplates and pooled for purification using nickel-sepharose columns (HisTrap HP, GE Healthcare, Chicago, IL, USA) and an AKTAxpress chromatography system (GE Healthcare). Briefly, nickel-charged columns were pre-eluted with elution buffer (10 mM Na_2_HPO_4_, 10 mM NaH_2_PO_4_, 0.5 M NaCl, 0.4 M imidazole, pH 7.4, filtered and degassed under vacuum) then equilibrated with running buffer (10 mM Na_2_HPO_4_, 10 mM NaH_2_PO_4_, 0.5 M NaCl, 0.04 M imidazole, pH 7.4, filtered and degassed under vacuum). Pooled supernatants were adjusted to approximately 0.1 M NaCl and 0.01 M imidazole then run through the column at a flow rate of 1 mL/min. The column was washed with 15 volumes of running buffer and then eluted in 0.5 mL fractions with 10 columns of elution buffer. Peak fractions were pooled and dialysed against PBS, then quantified by absorption at 280 nm using a Nanodrop 1000 instrument (Thermo Fisher Scientific).

Proteins used in surface plasmon resonance and crystallography were purified from conditioned medium collected from stable cell line cultures. The media was buffer exchanged with PBS and His-tagged proteins were captured with 5 mL HisTrap Excel columns (GE Healthcare), washed with 20 mM imidazole and eluted with 300 mM imidazole containing PBS buffer. The eluted proteins were further purified using a Superdex 200 16/60 column (GE Healthcare), in a buffer of 10 mM HEPES, pH 7.4, 150 mM NaCl. Before crystallisation, purified glycoproteins were deglycosylated using EndoF1.

### Antibody generation and purification

To characterise the expression pattern of the receptor protein, we raised and affinity purified a polyclonal antibody against the recombinant extracellular domain of zebrafish Cachd1.

Briefly, purified zebrafish Cachd1 ectodomain was prepared (see above) and sent to Cambridge Research Biochemicals (Billingham, United Kingdom) for a rabbit immunisation protocol. Activity against the Cachd1 ectodomain in rabbit blood sera was confirmed by ELISA. The blood serum was then affinity purified against biotinylated recombinant ectodomain immobilised on a streptavidin sepharose column, using an AKTAxpress chromatography system. Purified antibodies were eluted in fractions using a low pH buffer, then immediately neutralised. Peak fractions were tested for anti-Cachd1 activity, then pooled and dialysed against PBS. Total protein concentration was determined by absorbance at 280 nm by Nanodrop. The affinity purified antibody was checked for purity by SDS-PAGE and then validated by western blot, immunohistochemistry and flow cytometry (see Fig. S1 and data not shown) (*30*).

### Retrogenix Cell Microarray Technology

Cell Microarray Technology (*55*) was used to identify potential binding partners for multimerised human CACHD1 ectodomain (prepared as above) and was performed by Charles River Discovery Research Services UK Limited (formerly Retrogenix Limited; Chinley, United Kingdom; for bait target details, see (*56*)).

### Flow cytometry

To test Cachd1 prey binding interactions, live suspension culture cells transfected with *FZD-eGFP* constructs (and mock transfected control cells) were split into samples of 2.5-5.0 × 10^5^ cells in 1% BSA in PBS and placed in individual wells of 96 well round bottomed culture plates on ice. The cells were collected by centrifugation (200 × *g* for 5 minutes at 4°C) and then resuspended in dilutions of prey protein (human or zebrafish Cachd1 and mouse CD200R; batchwise dilution determined by β-lactamase assay, see above) or 1% BSA in PBS (secondary only controls), and incubated on ice for 30 minutes. The cells were washed three times, by centrifugation and resuspension in PBS, then incubated in anti-FLAG-phycoerythrin secondary antibody diluted in 1% BSA in PBS (Antibody registry ID: AB_1268475, mouse IgG1, 1:500, Abcam) for a further 30 minutes. Cells were washed three times, by centrifugation and resuspension in PBS, then analysed using a LSRFortessa flow cytometer with a 5-decade logarithmic scale for detection, a high throughput sampler for 96 well plates and FACSDiva software (BDBiosciences).

Where possible, we verified the surface expression of bait proteins or GFP-tagged Cachd1 in live cultures by following the same protocol but using specific primary antibodies in place of prey proteins (Fig. S1A, S11E; anti-Cachd1, diluted 1:700, bespoke rabbit polyclonal, see above; OMP-18R5, human anti-FZD7 IgG (*32*), diluted 1:2000, OncoMed Pharmaceuticals; anti-Smo, AB_1270802, rabbit polyclonal, diluted 1:500, Abcam; anti-Jamb, bespoke goat polyclonal, diluted 1:200, Everest Biotech) and Alexa Fluor-conjugated secondary antibodies (Molecular Probes, diluted 1:500).

The same procedure was followed for experiments testing the ability of OMP-18R5 to block Cachd1 prey-FZD-eGFP interactions, but with an additional incubation step before the application of prey proteins: cells were resuspended in OMP-18R5 diluted in 1% BSA in PBS (1:800) or 1% BSA in PBS only (control) and incubated for 30 minutes on ice, washed three times in PBS, and then resuspended in prey protein dilutions.

Mock transfection controls were used to determine forward and side scatter voltages for samples prior to data collection, and for background gating thresholds in data analysis. “Cells only” (no prey/primary or secondary antibodies) and “secondary antibody only” controls were included in every experiment. Flow cytometry data was analysed using FlowJo V10 (FlowJo, Ashland, OR, USA). Single cell populations were isolated using forward and side scatter values, bisected into eGFP-negative (untransfected) and eGFP-positive subpopulations and then the median value for phycoethryin fluorescence (indicating prey binding) calculated for each (Fig. S11B). Binding of prey protein to eGFP-positive cells was quantified by taking the ratio of the medians: 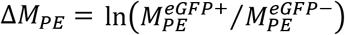.

### Surface Plasmon Resonance

Biotinylated proteins (FZD5/7/8CRD and LRP6P1E1P2E2 or LRP6P3E3P4E4) were obtained by co-transfection of avi3 tagged constructs (*57*) and a BirA-ER plasmid into HEK293T cells. About 500-1,000 resonance units of each of the biotinylated proteins were immobilized on a SA sensor chip (GE Healthcare), using a Biacore S200 machine (GE Healthcare) at 25 °C with a running buffer comprising 10 mM HEPES, pH 7.5, 150 mM NaCl and 0.005% Tween 20. A dilution series of purified CACHD1_ECD_ analyte was passed over the flow cells at high flow rate (100 μL/min) and the real-time response recorded at a frequency of 10 Hz. The response was plotted versus the concentration of the analyte and fitted by nonlinear regression to a one-site saturation binding model (Sigma Plot, Systat software, Inc. San Jose, CA).

### Crystallization, data collection and structure determination

Cachd1_ECD_ was concentrated to 5 mg ml^-1^, mixed with equal molar of FZD5_CRD_ and LRP6_P3E3P4E4_. The crystallization screening was carried out using the sitting-drop vapour diffusion method in 96-well plates. The crystals were obtained in condition of 0.1 M Calcium acetate; 0.1M Sodium acetate, pH 4.5; 10% (w/v) PEG 4000.

Crystals were flash frozen by immersion in a reservoir solution supplemented with 25% (v/v) glycerol followed by transfer to liquid nitrogen, and kept at −173 °C during X-ray data collection at I03, Diamond Light Source, with a wavelength of 0.9762 Å. The best diffracted crystal shows resolution of 4.7 Å, with space groups of *C*2_1_. Structure determination by molecular replacement with components structures solved in our laboratory and refinement used PHENIX (*58*) to good R factors and bond angles (see Table S2 for data collection and refinement statistics).

### *In situ* hybridization and Immunohistochemistry

Embryos or larvae were fixed in 4% paraformaldehyde and *in situs* performed following standard protocols (*59*). To create plasmid templates for *in situ* probe generation, regions of the *zgc:101731, slc18a3b, aoc1* and *cachd1* genes were PCR-amplified (see Table S5 for primer sequences) and TA-cloned into the pCRII vector. The *kiss1 in situ* probe template was generated directly by PCR (see Table S5). Previously published *in situ* probes used include (see Table S7 for references): *kctd12.1, kctd12.2, kctd8, axin2, lft1, otx5, spaw, selenop2, prss1, aldh1a3, dbx1b, wnt3a*, *lef1*. All enzymes used for plasmid linearization and *in vitro* transcription are listed in Table S6. Antisense probes were generated with digoxigenin and fluorescein labelling kits (Roche). Anti-digoxigenin-AP and anti-fluorescein-AP antibodies (Roche) coupled with 5-bromo-4-chloro-3’-indolyphosphate and nitro-blue tetrazolium chloride were used to visualize colorimetric *in situs*. Anti-digoxigenin-POD and anti-fluorescein-POD antibodies and Alexa Fluor-conjugated tyramides (Molecular Probes) were utilized for detection in fluorescent *in situ* hybridization.

*In situ* hybridisation chain reaction was performed according to published protocol (*60*) using Alexa Fluor-conjugated hairpin amplifiers and hybridisation buffers from Molecular Instruments Inc. (Los Angeles, CA, USA). Probe sets for *cachd1* and *lrp6* are detailed in Table S8.

For immunohistochemistry, embryos were stained according to published protocol, with the exception of using freshly fixed embryos without storage in methanol (*61*). Antibodies used in this study were: anti-acetylated α-tubulin (Antibody registry ID: AB_477585, clone 6-11B-1, mouse IgG2b, Sigma, diluted 1:250 in blocking solution), anti-SV2 (AB_2315387, mouse IgG1, deposited to the Developmental Studies Hybridoma Bank by Buckley, K.M., diluted 1:250), anti-HuC/HuD (AB_221448, clone 16A11, mouse IgG2b, Molecular Probes, diluted 1:250), anti-phospho-S10-histone H3 (AB_443110, mouse IgG1, Abcam, diluted 1:250), anti-Cachd1 (this study, rabbit polyclonal, diluted 1:50), anti-GFP (AB_10013661, rabbit polyclonal, Torrey Pines Biolab, diluted 1:1000; or AB_300798, chicken polyclonal, Abcam, diluted 1:500). The use of anti-PCNA (AB_2160343, clone PC10, mouse IgG2a, Cell Signalling Technology, diluted 1:100) required heat-mediated antigen retrieval: embryos were incubated in 10 mM Sodium citrate in PBS, pH 6.0, at 85°C for 20 mins before blocking. Alexa Fluor-conjugated anti-mouse IgG subtypes/rabbit/chicken secondary antibodies (Molecular Probes) were diluted 1:200 in blocking solution before use. 4’,6-diamidino-2-phenylindole (DAPI, Invitrogen) was added to embryos (10 μg/mL in PBST) to counterstain nuclei before imaging.

### Heat shock, laser cell ablation, BrdU, labelling of habenular projections and transplantation experiments

For rescue experiments, embryos transgenic for *Tg(HSE:cachd1,GFP)w160* were heat shocked for 30 minutes in a 40°C water bath, then raised at standard temperature to 4 dpf and fixed in 4% paraformaldehyde.

Laser cell ablation, BrdU incorporation experiments and lipophilic dye labelling of habenular efferent projections were performed as previously described (*16*).

### Imaging

For transmitted light pictures, larvae were mounted in glycerol and imaged using differential interference contrast optics (Leica CTR6000; 20× and 40× objectives; Leica Microsystems, Wetzlar, Germany). For confocal microscopy, heads were mounted in 1.2% low-melt agarose in glass-bottom dishes (MatTek, Ashland, MA, USA or LabTek, Grand Rapids, MI USA). Fluorescence was imaged by confocal laser scanning microscopy (Leica TCS SP5 and Leica TCS SP8) using a 40× oil-immersion objective (40× 1.3 Oil DIC III) or a 25× water-immersion objective (25× 0.95), and z stacks were acquired in 0.75 – 2 μm intervals. Cell cultures were imaged using an Marianis Spinning Disk (Intelligent Imaging Innovations, Inc., Denver, CO USA) system. 3D reconstructions and maximum-intensity projections were generated from stacks of images with Volocity (Improvision, Coventry, UK) and ImageJ (NIH). Image segmentation and quantification was performed using IMARIS (v8.0.1, Bitplane, Zurich, Switzerland).

### Organoid culture

Organoids were established from freshly isolated wild type small intestine or adenomas isolated from *Apc^min^* mice. Tissues were incubated in cold PBS containing 2 mM EDTA for isolating epithelial crypts and then cultured as described in (*62*), except that Matrigel was replaced with Cultrex© BME, Type 2 RGF PathClear (Amsbio, Abbingdon, UK, 3533-010-02). Briefly, organoid basal media contains EGF (Invitrogen, Waltham, MA, USA PMG8043), Noggin and R-spondin-1 (ENR). Noggin and R-spondin-1 conditioned media (CM) were generated from HEK293T cells. Wnt3a CM was generated from L cells.

### Quantitative RT-PCR

RNA was extracted from cell culture or organoids according to the manufacturer’s instructions (Qiagen RNeasy; Qiagen). cDNA was prepared using Maxima first strand cDNA synthesis kit with dsDNase (#1672, Thermo Fisher Scientific). Quantitative PCR detection was performed using PowerUp SYBR Green Master Mix (A25742, Applied Biosystems, Waltham, MA, USA). Assays for each sample were done in triplicate and were normalized to housekeeping genes *ACTB* (human *β-ACTIN*) or *Hrpt1* (mouse). Primer sequences are listed in the Table S5.

### Statistics

Statistical analysis was performed using RStudio (v1.4.1106, base R x64 v4.0.5, DescTools package v0.99.44) and Microsoft Excel 2010. Charts were plotted using the ggplot2 package (v3.3.3).

Descriptive statistics, scatterplots and normalised histograms for flow cytometry experiments were generated in FlowJo V10 (FlowJo, Ashland, OR, USA).

The Q’ test for equal proportions and modified Marascuilo procedure for multiple testing (using a Wilson variance calculation) are described in (*63*). Where the proportion was 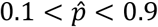 and/or *n* > 20, confidence intervals were calculated using a normal assumption; otherwise by the Wilson count method.

### Figure and manuscript preparation

Figures were compiled using Adobe Photoshop CS6 (64 bit).

Fluorescent and confocal microscopy images were adjusted globally for brightness and contrast using FIJI (v1.53n), scale bars added and then flattened into RGB images and exported as TIFFs.

Colour balance of wholemount *in situ* hybridisation images was adjusted in Adobe Photoshop CS6 (64 bit).

## References

1. M. L. Concha, I. H. Bianco, S. W. Wilson, Encoding asymmetry within neural circuits. Nat Rev Neurosci 13, 832–843 (2012).

2. L. J. Rogers, G. Vallortigara, Brain and behavioural asymmetries in non-human species. Laterality 26, v–vii (2021).

3. M. L. Concha, S. W. Wilson, Asymmetry in the epithalamus of vertebrates. J Anat 199, 63–84 (2001).

4. I. H. Bianco, S. W. Wilson, The habenular nuclei: a conserved asymmetric relay station in the vertebrate brain. Philos Trans R Soc Lond B Biol Sci 364, 1005–1020 (2009).

5. J. T. Gamse, C. Thisse, B. Thisse, M. E. Halpern, The parapineal mediates left-right asymmetry in the zebrafish diencephalon. Development 130, 1059–1068 (2003).

6. H. Aizawa et al., Laterotopic representation of left-right information onto the dorso-ventral axis of a zebrafish midbrain target nucleus. Curr Biol 15, 238–243 (2005).

7. I. H. Bianco, M. Carl, C. Russell, J. D. Clarke, S. W. Wilson, Brain asymmetry is encoded at the level of axon terminal morphology. Neural Dev 3, 9 (2008).

8. E. Dreosti, N. Vendrell Llopis, M. Carl, E. Yaksi, S. W. Wilson, Left-right asymmetry is required for the habenulae to respond to both visual and olfactory stimuli. Curr Biol 24, 440–445 (2014).

9. K. J. Turner et al., Afferent Connectivity of the Zebrafish Habenulae. Front Neural Circuits 10, 30 (2016).

10. N. Miyasaka et al., From the olfactory bulb to higher brain centers: genetic visualization of secondary olfactory pathways in zebrafish. J Neurosci 29, 4756–4767 (2009).

11. M. L. Concha et al., Local tissue interactions across the dorsal midline of the forebrain establish CNS laterality. Neuron 39, 423–438 (2003).

12. V. Duboc, P. Dufourcq, P. Blader, M. Roussigne, Asymmetry of the Brain: Development and Implications. Annu Rev Genet 49, 647–672 (2015).

13. S. Roberson, M. E. Halpern, Convergence of signaling pathways underlying habenular formation and axonal outgrowth in zebrafish. Development 144, 2652–2662 (2017).

14. M. Roussigne, P. Blader, S. W. Wilson, Breaking symmetry: the zebrafish as a model for understanding left-right asymmetry in the developing brain. Dev Neurobiol 72, 269–281 (2012).

15. M. Carl et al., Wnt/Axin1/beta-catenin signaling regulates asymmetric nodal activation, elaboration, and concordance of CNS asymmetries. Neuron 55, 393–405 (2007).

16. U. Husken et al., Tcf7l2 is required for left-right asymmetric differentiation of habenular neurons. Curr Biol 24, 2217–2227 (2014).

17. Y. S. Kuan et al., Distinct requirements for Wntless in habenular development. Dev Biol 406, 117–128 (2015).

18. L. Guglielmi et al., Temporal control of Wnt signaling is required for habenular neuron diversity and brain asymmetry. Development 147, dev182865 (2020).

19. R. Nusse, H. Clevers, Wnt/beta-Catenin Signaling, Disease, and Emerging Therapeutic Modalities. Cell 169, 985–999 (2017).

20. Z. Steinhart, S. Angers, Wnt signaling in development and tissue homeostasis. Development 145, dev146589 (2018).

21. D. Brafman, K. Willert, Wnt/beta-catenin signaling during early vertebrate neural development. Dev Neurobiol 77, 1239–1259 (2017).

22. E. Y. Rim, H. Clevers, R. Nusse, The Wnt Pathway: From Signaling Mechanisms to Synthetic Modulators. Annu Rev Biochem 91, 571–598 (2022).

23. B. J. Dean, B. Erdogan, J. T. Gamse, S. Y. Wu, Dbx1b defines the dorsal habenular progenitor domain in the zebrafish epithalamus. Neural Dev 9, 20 (2014).

24. H. Aizawa, M. Goto, T. Sato, H. Okamoto, Temporally regulated asymmetric neurogenesis causes left-right difference in the zebrafish habenular structures. Dev Cell 12, 87–98 (2007).

25. M. L. Concha, R. D. Burdine, C. Russell, A. F. Schier, S. W. Wilson, A nodal signaling pathway regulates the laterality of neuroanatomical asymmetries in the zebrafish forebrain. Neuron 28, 399–409 (2000).

26. M. Roussigne, I. H. Bianco, S. W. Wilson, P. Blader, Nodal signalling imposes left-right asymmetry upon neurogenesis in the habenular nuclei. Development 136, 1549–1557 (2009).

27. A. F. Schier, Nodal signaling in vertebrate development. Annu Rev Cell Dev Biol 19, 589–621 (2003).

28. J. T. Gamse et al., Directional asymmetry of the zebrafish epithalamus guides dorsoventral innervation of the midbrain target. Development 132, 4869–4881 (2005).

29. I. Lekk et al., Sox1a mediates the ability of the parapineal to impart habenular left-right asymmetry. Elife 8, e47376 (2019).

30. S. Dahimene et al., The alpha2delta-like Protein Cachd1 Increases N-type Calcium Currents and Cell Surface Expression and Competes with alpha2delta-1. Cell Rep 25, 1610–1621 e1615 (2018).

31. G. S. Cottrell et al., CACHD1 is an alpha2delta-Like Protein That Modulates CaV3 Voltage-Gated Calcium Channel Activity. J Neurosci 38, 9186–9201 (2018).

32. A. Gurney et al., Wnt pathway inhibition via the targeting of Frizzled receptors results in decreased growth and tumorigenicity of human tumors. Proc Natl Acad Sci U S A 109, 11717–11722 (2012).

33. S. Chen et al., Structural and functional studies of LRP6 ectodomain reveal a platform for Wnt signaling. Dev Cell 21, 848–861 (2011).

34. J. Wu et al., Structure of the voltage-gated calcium channel Ca(v)1.1 at 3.6 A resolution. Nature 537, 191–196 (2016).

35. L. Holm, Using Dali for Protein Structure Comparison. Methods Mol Biol 2112, 29–42 (2020).

36. H. Hirai, K. Matoba, E. Mihara, T. Arimori, J. Takagi, Crystal structure of a mammalian Wnt-frizzled complex. Nat Struct Mol Biol 26, 372–379 (2019).

37. C. Y. Janda, D. Waghray, A. M. Levin, C. Thomas, K. C. Garcia, Structural basis of Wnt recognition by Frizzled. Science 337, 59–64 (2012).

38. E. Bourhis et al., Reconstitution of a frizzled8.Wnt3a.LRP6 signaling complex reveals multiple Wnt and Dkk1 binding sites on LRP6. J Biol Chem 285, 9172–9179 (2010).

39. V. E. Ahn et al., Structural basis of Wnt signaling inhibition by Dickkopf binding to LRP5/6. Dev Cell 21, 862–873 (2011).

40. Z. Cheng et al., Crystal structures of the extracellular domain of LRP6 and its complex with DKK1. Nat Struct Mol Biol 18, 1204–1210 (2011).

41. M. Zebisch, V. A. Jackson, Y. Zhao, E. Y. Jones, Structure of the Dual-Mode Wnt Regulator Kremen1 and Insight into Ternary Complex Formation with LRP6 and Dickkopf. Structure 24, 1599–1605 (2016).

42. L. Novellasdemunt et al., USP7 Is a Tumor-Specific WNT Activator for APC-Mutated Colorectal Cancer by Mediating beta-Catenin Deubiquitination. Cell Rep 21, 612–627 (2017).

43. E. A. Rutledge, J. D. Benazet, A. P. McMahon, Cellular heterogeneity in the ureteric progenitor niche and distinct profiles of branching morphogenesis in organ development. Development 144, 3177–3188 (2017).

44. C. Y. Janda et al., Surrogate Wnt agonists that phenocopy canonical Wnt and beta-catenin signalling. Nature 545, 234–237 (2017).

45. F. J. van Eeden, M. Granato, J. Odenthal, P. Haffter, Developmental mutant screens in the zebrafish. Methods Cell Biol 60, 21–41 (1999).

46. R. N. Kettleborough et al., A systematic genome-wide analysis of zebrafish protein-coding gene function. Nature 496, 494–497 (2013).

47. J. C. Talbot, S. L. Amacher, A streamlined CRISPR pipeline to reliably generate zebrafish frameshifting alleles. Zebrafish 11, 583–585 (2014).

48. F. Kroll et al., A simple and effective F0 knockout method for rapid screening of behaviour and other complex phenotypes. Elife 10, e59683 (2021).

49. W. S. Talbot, A. F. Schier, Positional cloning of mutated zebrafish genes. Methods Cell Biol 60, 259–286 (1999).

50. K. M. Bushell, C. Sollner, B. Schuster-Boeckler, A. Bateman, G. J. Wright, Large-scale screening for novel low-affinity extracellular protein interactions. Genome Res 18, 622–630 (2008).

51. G. Lennon, C. Auffray, M. Polymeropoulos, M. B. Soares, The I.M.A.G.E. Consortium: an integrated molecular analysis of genomes and their expression. Genomics 33, 151–152 (1996).

52. C. Sollner, G. J. Wright, A cell surface interaction network of neural leucine-rich repeat receptors. Genome Biol 10, R99 (2009).

53. Y. Zhao, J. Ren, S. Padilla-Parra, E. E. Fry, D. I. Stuart, Lysosome sorting of beta-glucocerebrosidase by LIMP-2 is targeted by the mannose 6-phosphate receptor. Nat Commun 5, 4321 (2014).

54. P. J. Reeves, N. Callewaert, R. Contreras, H. G. Khorana, Structure and function in rhodopsin: high-level expression of rhodopsin with restricted and homogeneous N-glycosylation by a tetracycline-inducible N-acetylglucosaminyltransferase I-negative HEK293S stable mammalian cell line. Proc Natl Acad Sci U S A 99, 13419–13424 (2002).

55. J. Freeth, J. Soden, New Advances in Cell Microarray Technology to Expand Applications in Target Deconvolution and Off-Target Screening. SLAS Discov 25, 223–230 (2020).

56. L. Turner et al., Severe malaria is associated with parasite binding to endothelial protein C receptor. Nature 498, 502–505 (2013).

57. A. R. Aricescu, W. Lu, E. Y. Jones, A time- and cost-efficient system for high-level protein production in mammalian cells. Acta Crystallogr D Biol Crystallogr 62, 1243–1250 (2006).

58. T. C. Terwilliger et al., Iterative model building, structure refinement and density modification with the PHENIX AutoBuild wizard. Acta Crystallogr D Biol Crystallogr 64, 61–69 (2008).

59. C. Thisse, B. Thisse, High-resolution in situ hybridization to whole-mount zebrafish embryos. Nat Protoc 3, 59–69 (2008).

60. H. M. Choi et al., Mapping a multiplexed zoo of mRNA expression. Development 143, 3632–3637 (2016).

61. K. J. Turner, T. G. Bracewell, T. A. Hawkins, Anatomical dissection of zebrafish brain development. Methods Mol Biol 1082, 197–214 (2014).

62. T. Sato et al., Single Lgr5 stem cells build crypt-villus structures in vitro without a mesenchymal niche. Nature 459, 262–265 (2009).

63. G. A. Michael, A significance test of interaction in 2 x K designs with proportions. TQMP 3, 1–7 (2007).

64. B. Kilian et al., The role of Ppt/Wnt5 in regulating cell shape and movement during zebrafish gastrulation. Mech Dev 120, 467–476 (2003).

65. S. Witzel, V. Zimyanin, F. Carreira-Barbosa, M. Tada, C. P. Heisenberg, Wnt11 controls cell contact persistence by local accumulation of Frizzled 7 at the plasma membrane. J Cell Biol 175, 791–802 (2006).

66. Z. M. Varga et al., Zebrafish smoothened functions in ventral neural tube specification and axon tract formation. Development 128, 3497–3509 (2001).

67. G. Weidinger, C. J. Thorpe, K. Wuennenberg-Stapleton, J. Ngai, R. T. Moon, The Sp1-related transcription factors sp5 and sp5-like act downstream of Wnt/beta-catenin signaling in mesoderm and neuroectoderm patterning. Curr Biol 15, 489–500 (2005).

68. T. Kudoh et al., A gene expression screen in zebrafish embryogenesis. Genome Res 11, 1979–1987 (2001).

69. F. Biemar et al., Pancreas development in zebrafish: early dispersed appearance of endocrine hormone expressing cells and their convergence to form the definitive islet. Dev Biol 230, 189–203 (2001).

70. S. Long, N. Ahmad, M. Rebagliati, The zebrafish nodal-related gene southpaw is required for visceral and diencephalic left-right asymmetry. Development 130, 2303–2316 (2003).

71. C. Thisse, B. Thisse, Antivin, a novel and divergent member of the TGFbeta superfamily, negatively regulates mesoderm induction. Development 126, 229–240 (1999).

72. G. Lupo et al., Retinoic acid receptor signaling regulates choroid fissure closure through independent mechanisms in the ventral optic cup and periocular mesenchyme. Proc Natl Acad Sci U S A 108, 8698–8703 (2011).

73. S. Krauss, V. Korzh, A. Fjose, T. Johansen, Expression of four zebrafish wnt-related genes during embryogenesis. Development 116, 249–259 (1992).

74. L. E. Valdivia et al., Lef1-dependent Wnt/beta-catenin signalling drives the proliferative engine that maintains tissue homeostasis during lateral line development. Development 138, 3931–3941 (2011).

