## Supplementary Figures and Tables for "Cachd1 is a novel Frizzled- and LRP6-interacting protein required for neurons to acquire left-right asymmetric character"

**This file includes:**

Supplementary Figures S1-S15

Supplementary Tables S1-S8

### Supplementary figures

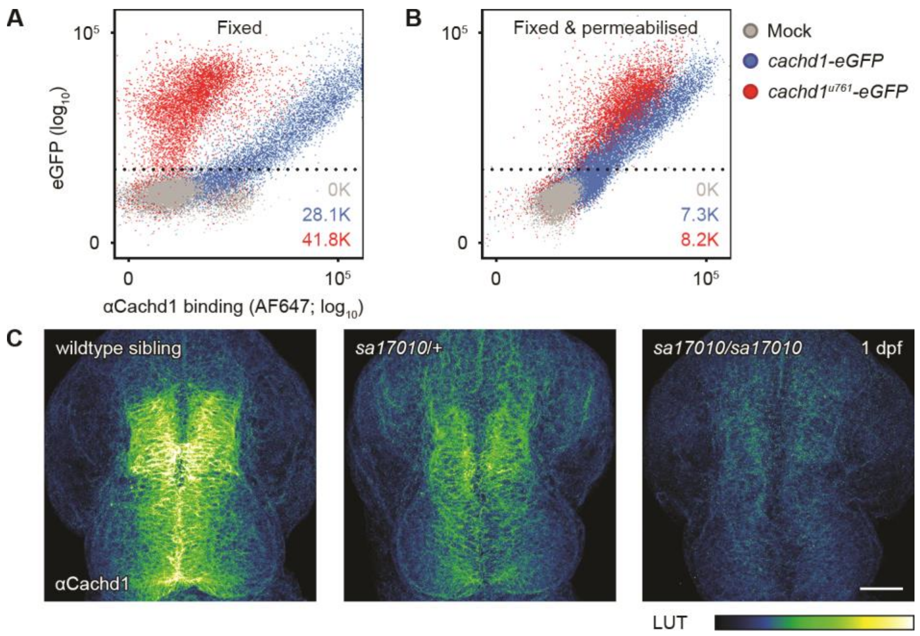


### Fig. S1. Cross-validation of *cachd1* mutations and anti-Cachd1 antibody.

(A and B) Flow cytometry of untransfected (Mock, grey) and transfected HEK293E cells expressing wildtype Cachd1- (blue) or mutant Cachd1$^{u761}$-eGFP (red) fusion proteins stained with anti-Cachd1 antibody after fixation (A) and permeabilization (B). The mutant fusion protein was stained strongly after permeabilization of the cells, suggesting the protein is not trafficked to the plasma membrane. Numbers in each panel indicate the total number of eGFP-positive events recorded.

(C) Dorsal views of brains of 1 dpf siblings from a *cachd1*$^{sa17010}$/+ incross, stained with anti-Cachd1 antibody. Intensity of staining of the midbrain roofplate and dorsal diencephalon is correlated to *sa17010* genotype (see Table S1), suggesting the early nonsense mutation prevents translation of the protein. Maximum projections of confocal stacks.

Scale bars = 50 µm.

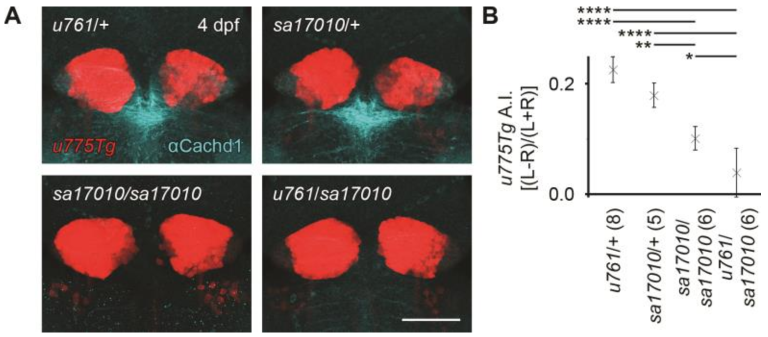


**Fig. S2. The *sa17010* allele of *cachd1* phenocopies, and is unable to complement, *u761*.**

(A) Dorsal views of 4 dpf *Tg*(*110316_GFP*)*u775/o* larvae carrying *u761* and/or *sa17010* alleles of *cachd1*, stained with anti-Cachd1 antibody (cyan). *Tg*(*110316_GFP*)*u775* express GFP in differentiated dorsal habenula neurons (red). Right habenula labelling is larger in the *sa17010* and *u761*/*sa17010* larvae than in heterozygotes, indicating bilateral 'double left' symmetry. Maximum projections of confocal stacks. Scale bar = 50 µm.

(B) Asymmetry index calculated from GFP habenulae volumes of *u761*/+, *sa17010*/+, *sa17010/sa17010* and *u761*/*sa17010* 4 dpf larvae. *sa17010* homozygotes and *u761*/*sa17010* transheterozyotes are bilaterally symmetric, suggesting loss of function of *cachd1* is causative for the *rorschach* phenotype. Number of larvae analysed indicated in brackets. Error bars represent 95% confidence intervals of the mean. ANOVA (degrees of freedom = 3, F = 27.41, $p = 1.87 \times 10^{-7}$) and *post hoc* Tukey pairwise comparisons was used for hypothesis testing, * $0.1 > p > 0.05$, ** $0.05 > p > 0.01$, **** $p < 0.005$.

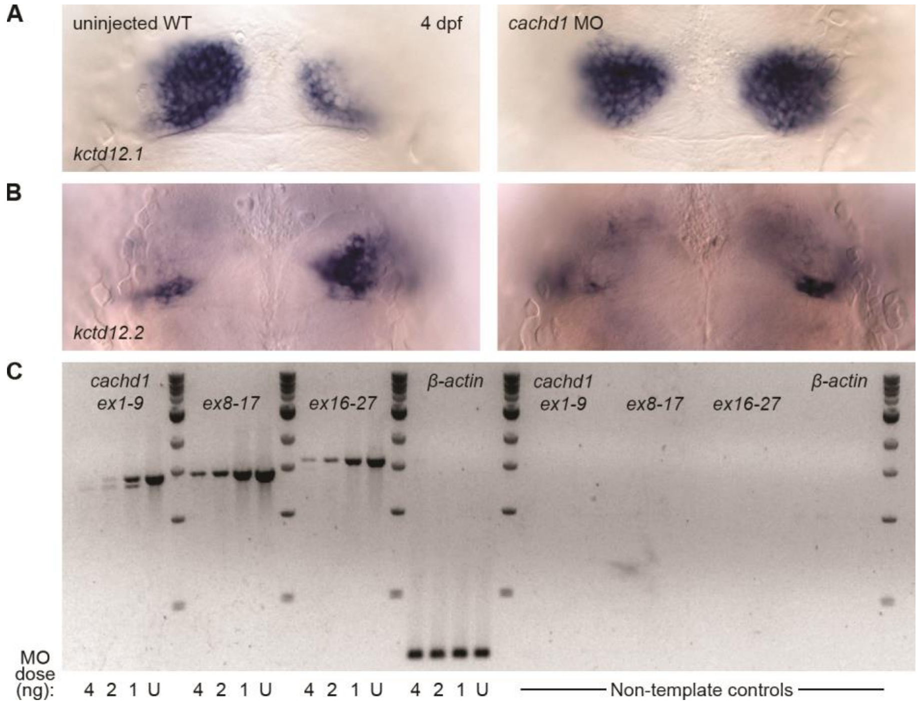


**Fig. S3. Morpholino knockdown of *cachd1* results in bilateral symmetry.**

(A-B) Dorsal views of 4 dpf uninjected wildtype and *cachd1* morpholino-injected larvae after wholemount *in situ* hybridisation using antisense riboprobes against asymmetric dorsal habenula markers *kctd12.1* (A, n = 253/263) or *kctd12.2* (B, n = 16/17). Note the increase in *kctd12.1* expression and corresponding decrease of *kctd12.2* expression in the right habenula of *cachd1* morphants.

(C) Semi-quantitative RT-PCR for *cachd1* transcripts (three primer sets spanning exons 1-9, exons 8-17 and exons 16-27) and reference gene *β–actin* in uninjected embryos (U) and those injected with ~4 ng, 2 ng and 1 ng of *cachd1* morpholino (MO1) showing a dose dependent reduction in *cachd1* expression and mis-splicing (exon 1-9). Subsequent Sanger sequencing of the RT-PCR products indicated mis-splicing resulted in an 89 bp deletion from the 3' end of exon 1 and usage of a cryptic donor site.

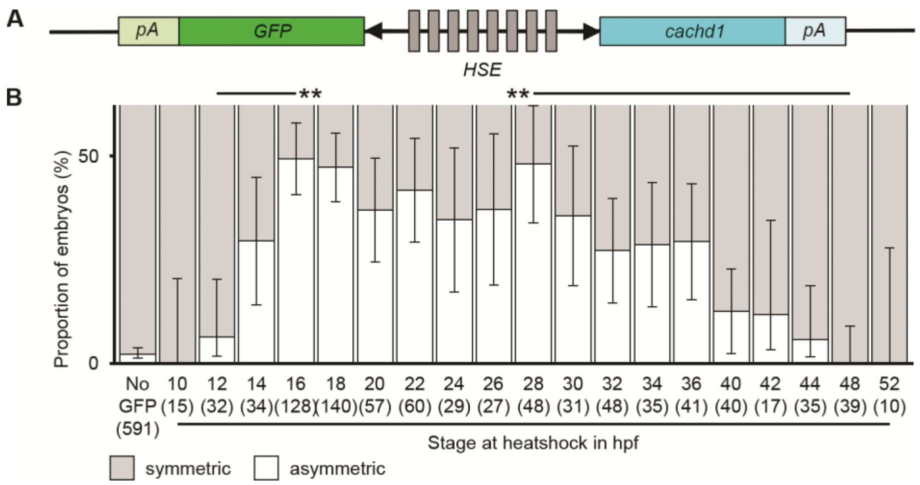


**Fig. S4. Asymmetry in *cachd1$^{u761}$* mutants is restored by overexpression of wildtype *cachd1*.**

(A) Schematic of construct used to generate transgenic *Tg(hse:cachd1*, *GFP)w160* fish. A bidirectional heat shock promoter (HSE) drives simultaneous expression of GFP and Cachd1 in response to acute exposure to heat.

(B) Chart showing the percentage of GFP-positive, *cachd1$^{u761}$* mutant larvae with symmetric (grey) or asymmetric (white) *kctd12.1* expression at 4 dpf, after receiving a heat shock at the stage indicated (GFP-negative, heat-shocked sibling larvae included in ‘No GFP’ column). There was significant restoration of asymmetry after heat shock between 16 h. p. f. - 28 h. p. f., after which the proportion of larvae with wildtype phenotype declined. Number of larvae indicated in brackets. Error bars represent 95% confidence interval for the proportion, calculated using a normal approximation, or Wilson score when the proportion was less than 0.1, and/or the number of larvae tested was less than 20. Q' test of equality of proportions (degrees of freedom = 19, $\chi^2$ = 342.27, p = 4.03 × $10^{-61}$) and *post hoc* modified Marascuilo procedure for multiple comparisons of proportions with Benjamini & Hochberg correction for multiple testing was used to test significance, ** 0.05 > p > 0.01. A limited number of statistically significant differences are presented here for clarity.

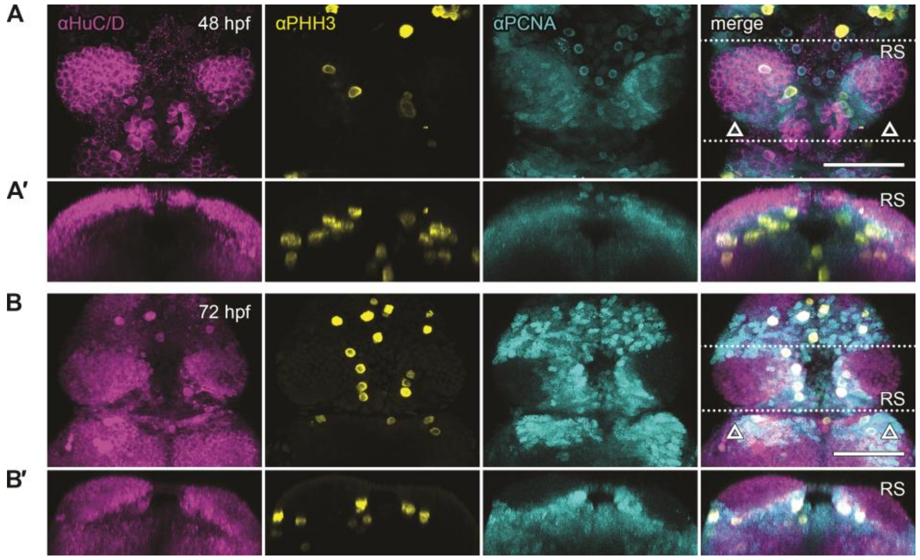


**Fig. S5. The periventricular domain ventral to the pineal complex is proliferative.**

(A-B′) Dorsal (A, B) and transverse projections (A′, B′) of 48 hpf (A, A′) and 72 hpf (B, B′) embryos stained with anti-HuC/D to mark differentiated neurons (magenta), anti-phospho-histone H3 to mark neuronal cells in M-phase (yellow) and anti-PCNA to mark neuronal cells in G1/S phase (cyan) of the cell cycle. Maximum projections of confocal stacks. The positions of habenulae are indicated with open arrowheads. The dotted lines in (A, B) indicate the volume shown in the transverse projections (A′, B′). Note that staining with anti-PCNA required antigen retrieval steps that inhibited anti-Cachd1 labelling, preventing co-staining; compare to Cachd1 expression in Fig. 1, 2, S1 and S2.

Scale bars = 50 µm.

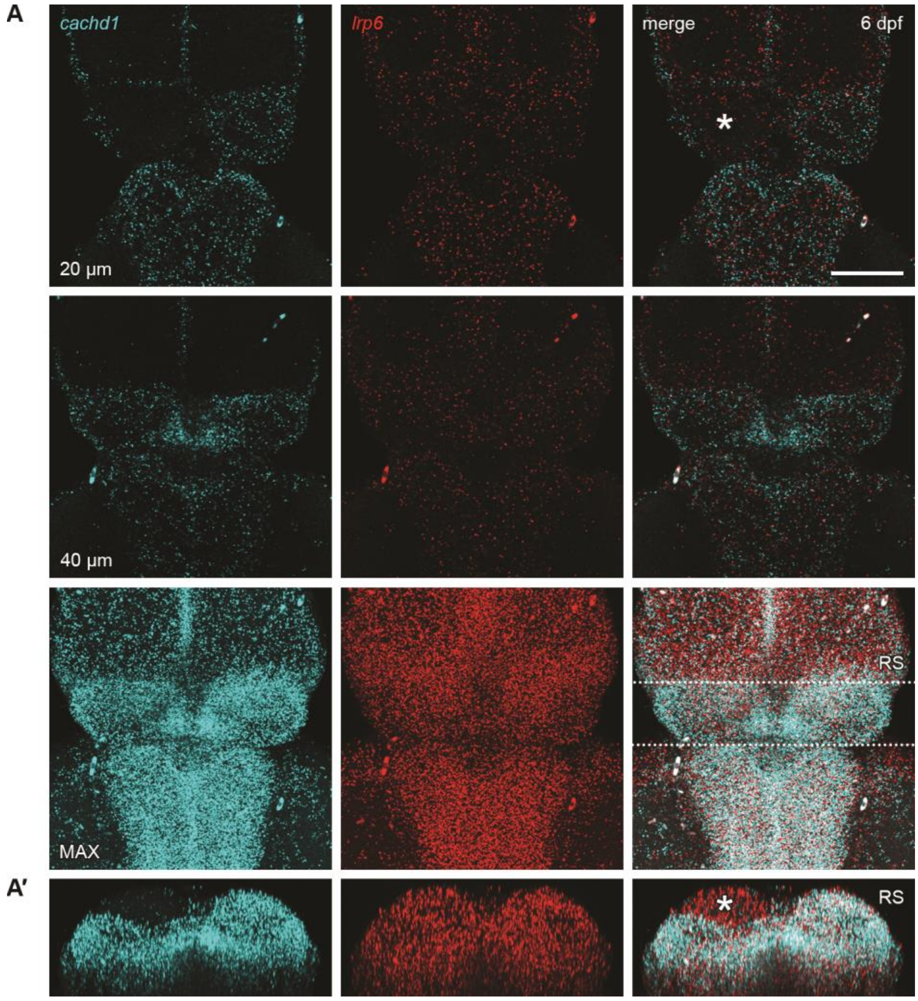


### <u>Fig. S6. Expression of *cachd1* is asymmetric at larval stages.</u>

(A-A′) Dorsal view (A) and transverse projection (A′) of a dissected 6 dpf wildtype larva after *in situ* hybridisation chain reaction with probes against *cachd1* (cyan) and *lrp6* (red).

(A) Single confocal slices at ~20 µm and 40 µm depth (from dorsal brain surface) and a maximum projection of the same confocal stack. Dotted lines indicate the approximate volume presented in the transverse projection (RS, A′). *cachd1* expression remains in the periventricular habenular domain ventral to the pineal, but is also expressed in the right habenula in its entirety; it is absent from the dorsalmost domain of the left dorsal habenula (asterisk). *lrp6* is expressed ubiquitously.

Scale bar = 50 µm.

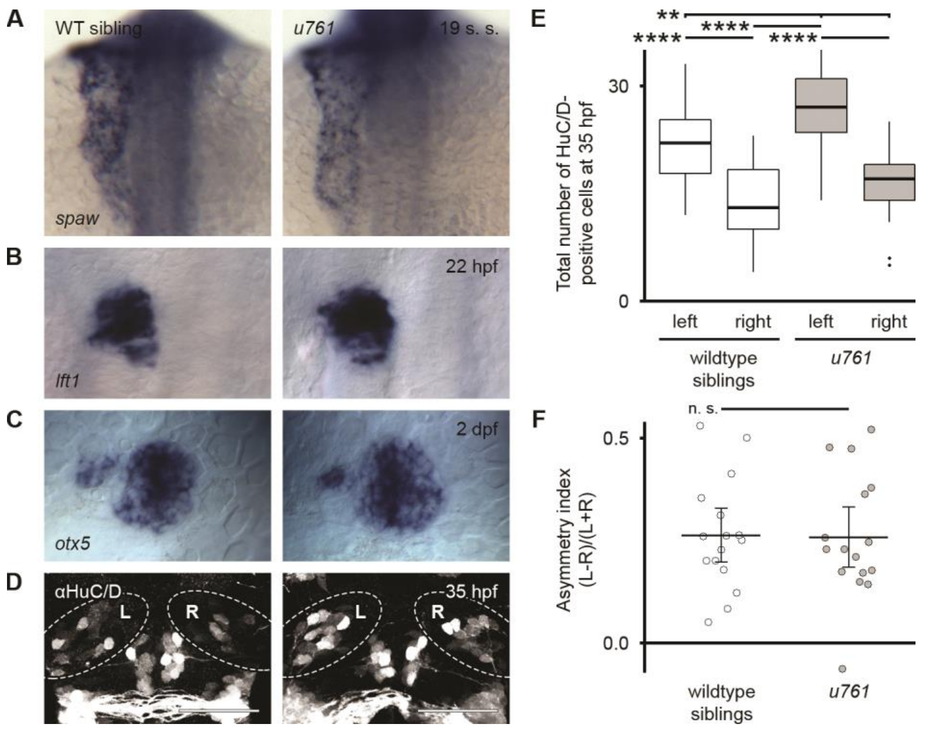


### Fig. S7. Early nodal-related left-right asymmetries are unperturbed in *cachd1*$^{u761}$ mutants.

(A-C) Dorsal views of wildtype sibling and *cachd1*$^{u761}$ mutant embryos after colorimetric wholemount *in situ* hybridisation with antisense riboprobes for the *nodal* signalling pathway ligand-encoding gene *spaw* (A, 19 somite stage) or target, *lft1* (B, 22 hpf), or pan-pineal complex marker *otx5* (C, ~2 dpf) indicating early asymmetric expression of these marker genes is unaffected in *cachd1*$^{u761}$ mutants.

(D) Dorsal views of wildtype or *cachd1*$^{u761}$ mutant embryos stained with anti-HuC/D to mark differentiated neurons at ~35 hpf. Left and right dorsal habenula indicated with dotted lines. Note the overall increase in the number of differentiated neurons in the left and right dorsal habenula of *cachd1*$^{u761}$ mutants (quantified in E). Scale bars = 50 µm.

(E) Boxplots showing quantification of the number of anti-HuC/D-positive nuclei in the left and right dorsal diencephalon of wildtype siblings (white) and *cachd1*$^{u761}$ mutants (grey); n = 16 for both groups. Kruskal-Wallis rank sum test (degrees of freedom = 3, $\chi^2$ = 27.208, p = 5.32 × $10^{-6}$) and *post hoc* pairwise comparisons using Wilcoxon rank sum test with continuity correction and Benjamini & Hochberg correction for multiple testing, ** 0.05 > p > 0.01, **** p < 0.005.

(F) Dotplots of the asymmetry index calculated for each wildtype sibling and *cachd1*$^{u761}$ mutant embryo, based on the number of anti-HuC/D-positive nuclei. Although there is an overall increase in early neurogenesis in *cachd1*$^{u761}$ mutants, there is a leftward bias, consistent with correct early Nodal-related asymmetry determination. Bar indicates sample mean and error bars indicate 95% confidence intervals of the mean. Welch two sample, two-tailed, *t*-test (degrees of freedom = 29.629, *t* = 0.094, p = 0.926).

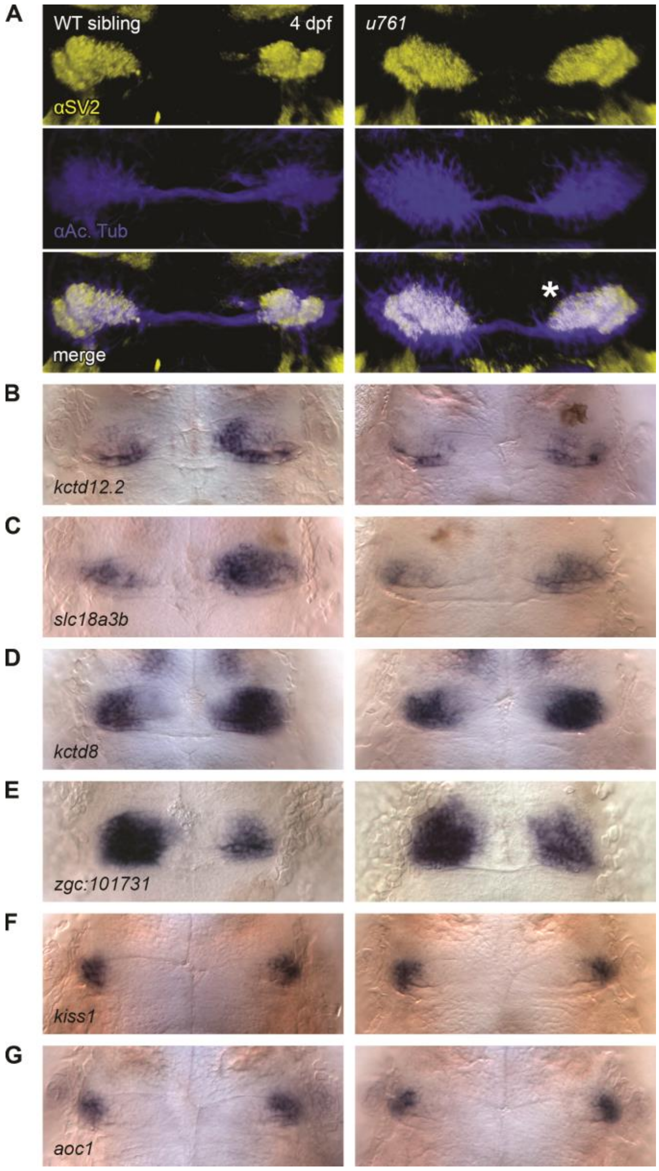


**Fig. S8. Symmetric neuroanatomy and gene expression in the dorsal habenulae of *cachd1*$^{u761}$ mutants.**

(A) Dorsal views of immunohistochemistry labelling neuropil (anti-SV2, yellow) and axons (anti-acetylated tubulin, blue) in the habenulae in 4 dpf wildtype and *cachd1*$^{u761}$ mutant larvae. Asterisk marks the increased volume of $dHb_L$-associated neuropil in the right habenula of *cachd1*$^{u761}$ mutants. Maximum projections of confocal stacks.

(B-G) Dorsal views of 4 dpf wildtype and *cachd1*$^{u761}$ larvae after colorimetric wholemount *in situ* hybridisation with riboprobes against dorsal habenula markers *kctd12.2* (B), *slc18a3b* (C), *kctd8* (D), *zgc:101731* (E) and ventral habenula markers *kiss1* (F) and *aoc1* (G). The asymmetric dorsal habenula markers are reduced or symmetric in *cachd1*$^{u761}$ mutants, but the ventral habenula markers are unaffected, suggesting *cachd1* does not play a role in neurogenesis of the ventral habenula.

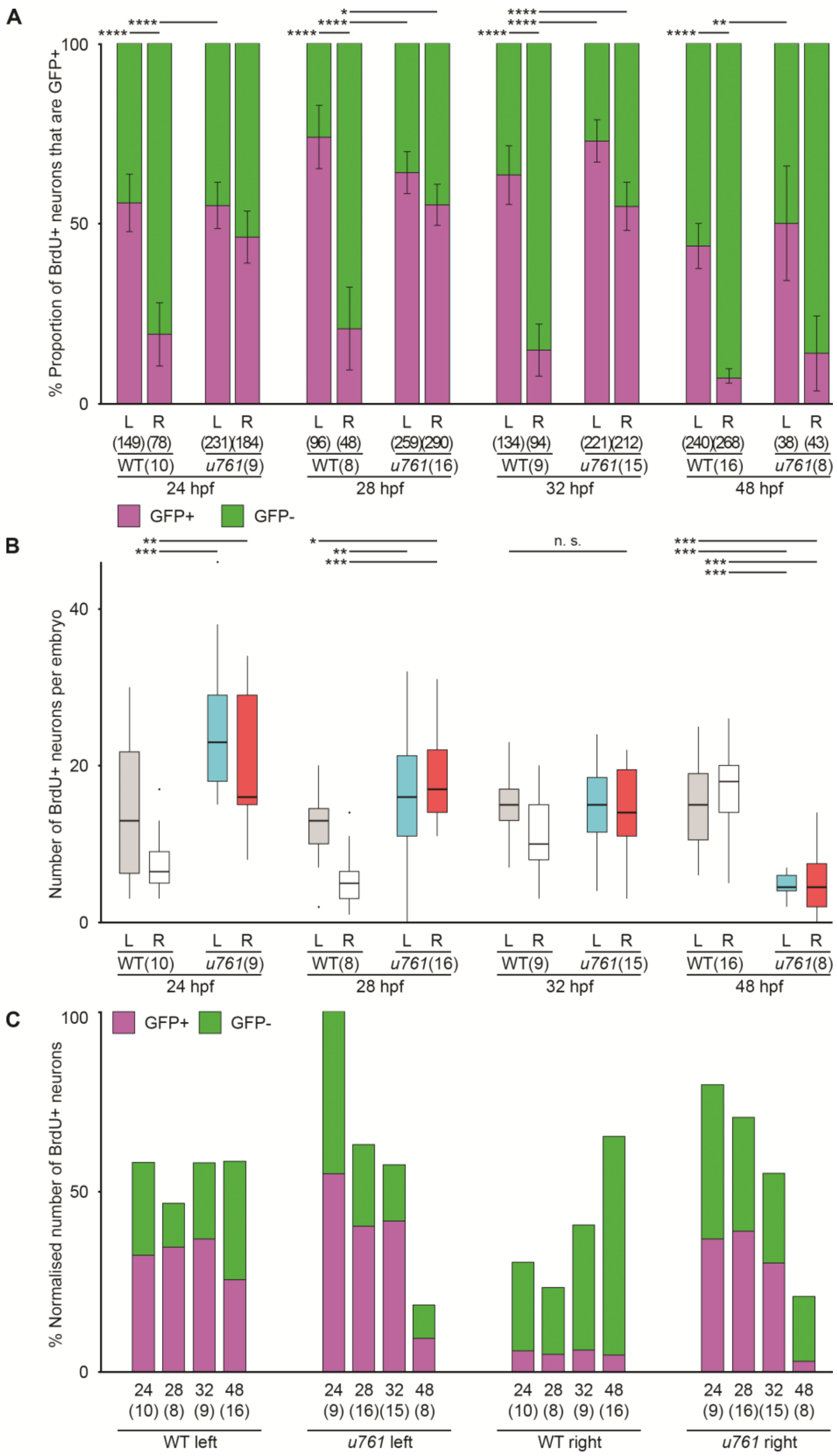


**Fig. S9. Precocious neurogenesis in *cachd1*$^{u761}$ mutant embryos with increased likelihood of acquiring lateral fate.**

(A) Bar chart showing the proportion of BrdU-positive cells that express the $dHb_L$ marker transgene *pku588Et* (GFP+, magenta) in ~5 dpf wildtype sibling and *cachd1*$^{u761}$ mutants given a pulse of BrdU at different stages (24 hpf – 48 hpf). Note the change in proportion in the right habenula of *cachd1*$^{u761}$ mutants, indicating an increased likelihood of acquiring $dHb_L$ character.

Error bars represent 95% confidence intervals of the proportion, calculated using a normal approximation, or Wilson score when the proportion was less than 0.1, and/or the number of larvae tested was less than 20. The total number of cells and larvae in each condition is indicated in brackets. Samples for 24 hpf are replicated from Fig. 3 for comparison. Q' test of equality of proportions (degrees of freedom = 15, $\chi^2$ = 747.49, p = 1.39 × $10^{-149}$) and *post hoc* pairwise comparisons using a modified Marascuilo procedure with Benjamini & Hochberg correction for multiple testing, * 0.1 > p > 0.05, ** 0.05 > p > 0.01, **** p < 0.005. A limited number of significant differences are presented here for clarity (within each BrdU pulse timepoint only).

(B) Boxplot showing the number of BrdU-positive cells in the habenulae of ~5 dpf wildtype sibling (grey left, white right) and *cachd1*$^{u761}$ mutants (cyan left, red right) given a pulse of BrdU at different stages (24 hpf – 48 hpf). Note the significant increase in neurogenesis at early stages and decrease at later stages in *cachd1*$^{u761}$ mutants. The number of larvae in each condition is indicated in brackets. Kruskal-Wallis rank sum test (degrees of freedom = 15, $\chi^2$ = 70.975, p = 2.99 × $10^{-9}$) and *post hoc* pairwise comparisons using Wilcoxon rank sum test with continuity correction and Benjamini & Hochberg correction for multiple testing, n. s. p > 0.1, * 0.1 > p > 0.05, ** 0.05 > p > 0.01, *** 0.01 > p > 0.005, **** p < 0.005. A limited number of significant differences are presented here for clarity (within each BrdU pulse timepoint only).

(C) Summary bar chart showing the total number of BrdU-positive neurons observed in the habenula of ~5 dpf in a wildtype and *cachd1*$^{u761}$ mutant larvae labelled at each BrdU pulse timepoint that were expressing the $dHb_L$ marker transgene *pku588Et* (GFP+, magenta) or not (GFP-, green), normalised to the highest number of observed labelled neurons and the number of larvae in each condition. Note for example, the increased proportion of GFP+ neurons born on the right hand side of *cachd1*$^{u761}$ mutants at early stages compared to WT (significant differences identified in A) and the increased total number of neurons (significant differences identified in B).

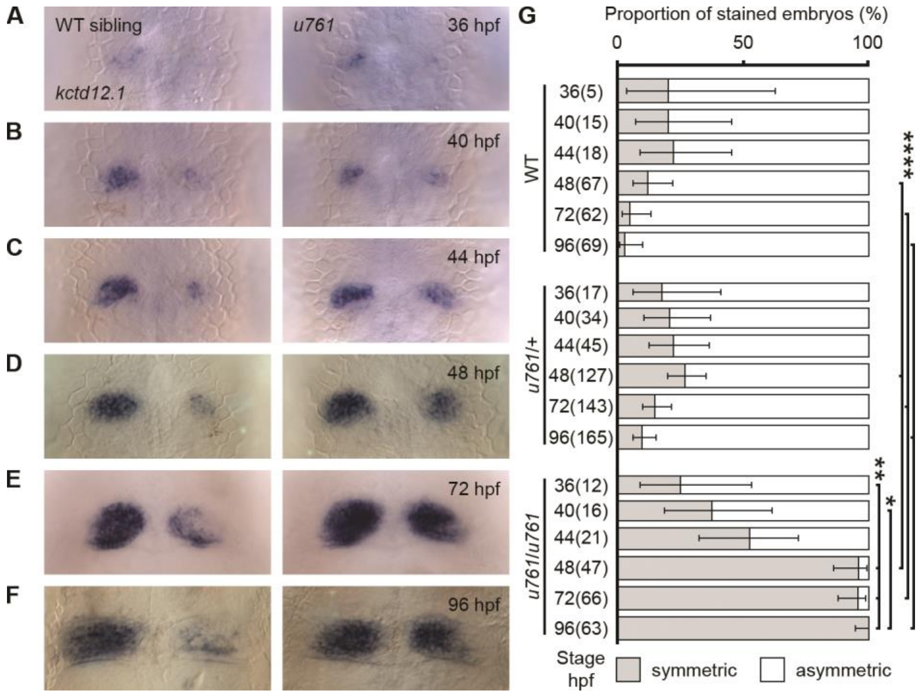


**Fig. S10. Biased acquisition of left-sided character in right dorsal habenula of *cachd1*$^{u761}$ mutants.**

(A-F) Dorsal views of wildtype sibling and *cachd1*$^{u761}$ mutant embryos at different developmental stages, 36 hpf (A) to 96 hpf (F), after colorimetric wholemount *in situ* hybridisation with an antisense riboprobe for the $dHb_L$ marker *kctd12.1*. Expression of *kctd12.1* is increased in the right habenula of *cachd1*$^{u761}$ mutants over the timecourse, consistent with an increased likelihood of acquiring $dHb_L$ fate.

(G) Bar chart showing the proportion of wildtype, *u761*/+ and *u761/u761* mutants embryos that showed symmetric (grey) or overtly asymmetric (white) *kctd12.1* staining at the different stages. Error bars represent the 95% confidence intervals for the proportion calculated using the Wilson score. Number of embryos in each condition is indicated in brackets. Q' test of equality of proportions (degrees of freedom = 17, $\chi^2$ = 646.41, p = 2.08 × $10^{-126}$) and *post hoc* modified Marascuilo procedure for multiple comparisons of proportions with Benjamini-Hochberg correction for multiple testing was used to test significance, ** 0.05 > p > 0.01, **** p < 0.005.

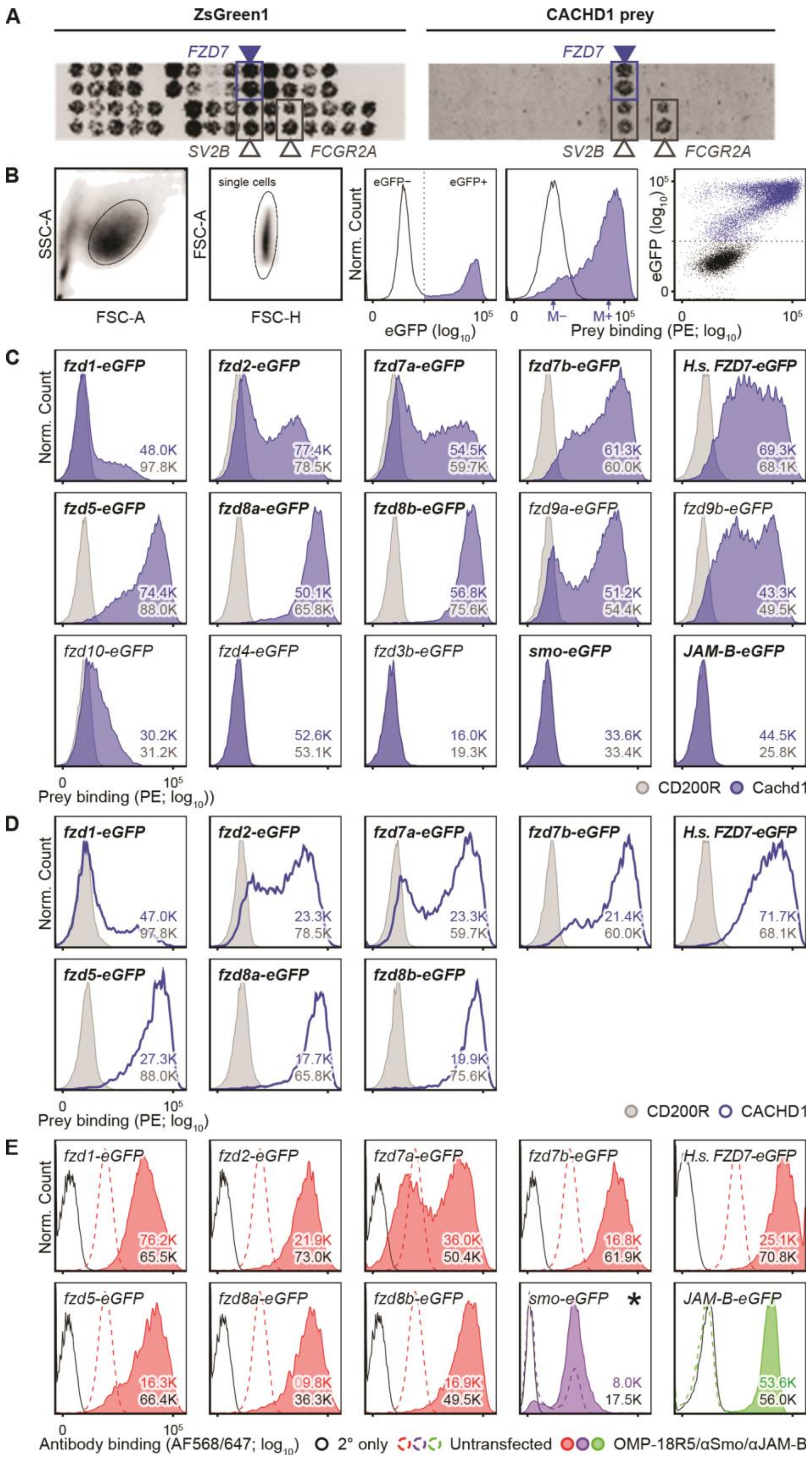


**Fig. S11. Physical interactions of Cachd1 with Fzd family proteins detected by cell microarray and flow cytometry.**

(A) Cell microarray data showing the interaction between a human CACHD1 prey protein and its target FZD7 (blue). Expression vectors encoding both ZsGreen1 and FZD7, together with a range of interactions from alternative prey proteins were spotted onto slides, and human HEK293 cells were reverse transfected. Fixed cells were subsequently incubated with human CACHD1 prey protein, and detected with an AlexaFluor-647 conjugated secondary antibody. ZsGreen fluorescence was used to confirm transfection efficiency and spot locations on the slides (left-hand panel). Two other hits, SV2B and FCGR2A, were considered false positives because of multiple binding interactions with a wide range of prey proteins.

(B) Gating strategy for testing specific interactions with transiently transfected cells. Single cells were isolated by forward (FSC-A, FSC-H) and side (FSC-A) light scatter, then separated into eGFP-negative (untransfected or not expressing the eGFP fusion protein bait; black outline) and eGFP-positive (transfected, blue) gates. Prey binding is indicated by increased PE fluorescence in the eGFP-positive population. The ratio of median PE fluorescence of either subpopulation ($\Delta M_{PE} = \ln\left(M_{PE}^{eGFP+}/M_{PE}^{eGFP-}\right)$) was used to quantify the degree of prey binding to the eGFP-positive population.

(C, D) Examples of normalized histograms of eGFP-positive populations for each eGFP fusion protein bait transfection indicated, tested with either zebrafish Cachd1 (C, solid blue), human CACHD1 (D, blue outline) or CD200R negative control prey (grey). Note that human CACHD1 prey is able to bind zebrafish Fzd family proteins and *vice versa*, suggesting conservation of binding (not all combinations tested). Numbers in each panel indicate the total number of eGFP-positive events collected for each condition over all replicates. Bold transfection titles indicate validation of surface expression with antibodies.

(E) Examples of normalized histograms of eGFP-positive populations for each eGFP fusion protein bait tested with antibodies to detect surface expression of Fzd family proteins (OMP-18R5, red), smo-eGFP (purple) or negative control bait protein JAM-B-eGFP (green). The dotted line in each panel indicates antibody binding fluorescence in mock transfected HEK293E cells; black outline indicates the secondary only negative control. Note that untransfected HEK293E cells appear to have endogenous surface expression of Fzd family receptor(s) and the Smo receptor (dotted lines). Numbers in each panel indicate the total number of eGFP-positive events collected for each condition. Asterisk indicates a formaldehyde fixed cell population.

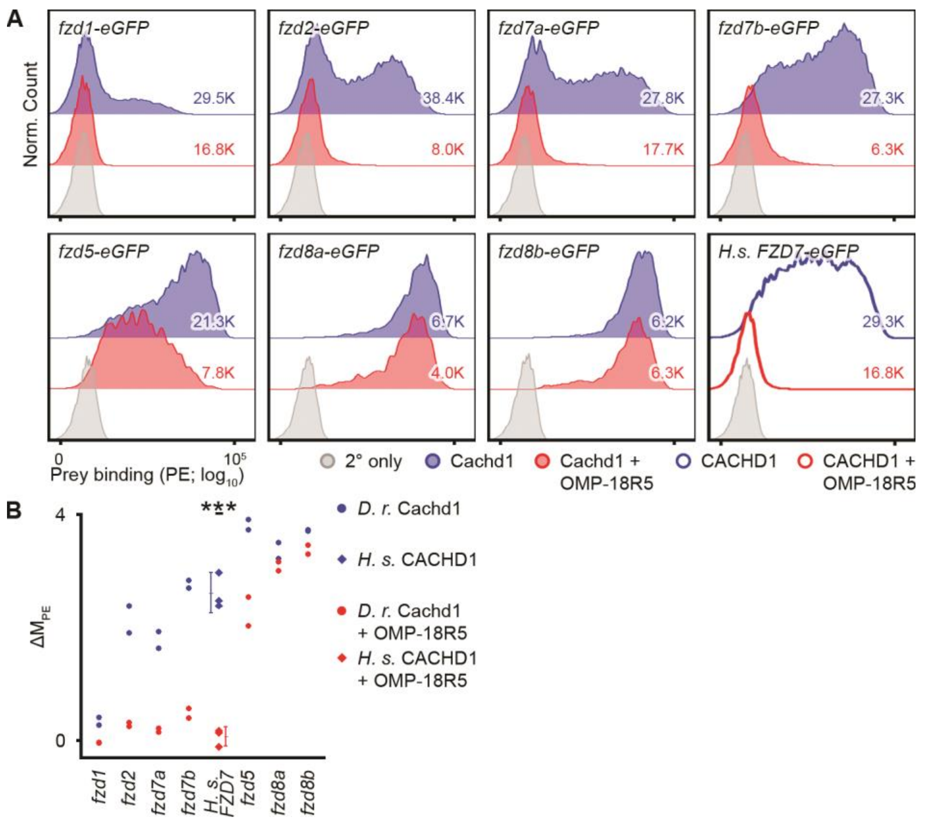


### Fig. S12. Cachd1 prey binding to Fzd family receptors blocked by anti-Fzd antibody.

(A) Examples of normalized histograms of eGFP-positive populations for each eGFP fusion protein bait transfection indicated, tested with either zebrafish Cachd1 prey protein alone (blue) or after pre-incubation of the cells with OMP-18R5 antibody (red). Human *FZD7-eGFP* histogram duplicated from Fig. 4A for comparison. Secondary antibody only control shown in grey. Total number of events collected for each condition indicated in each plot. Note that Cachd1 prey binding to zebrafish Fzd family bait proteins Fzd1/2/7a/7b is effectively blocked by OMP-18R5 pre-incubation, but not to Fzd5/8a/8b.

(B) Dotplot showing $\Delta M_{PE}$ for transfections tested for Cachd1 prey protein interaction (circles and diamonds represent zebrafish and human prey proteins respectively) with (red) or without pre-incubation with OMP-18R5 (blue). Mean indicated with a single line, error bars indicate 95% confidence intervals of the mean. One-tailed paired *t*-test for human CACHD1-FZD7 interaction only (degrees of freedom = 2, $t$ = 9.53, *** p = 0.0054), due to limited numbers of replicates for other inhibition tests.

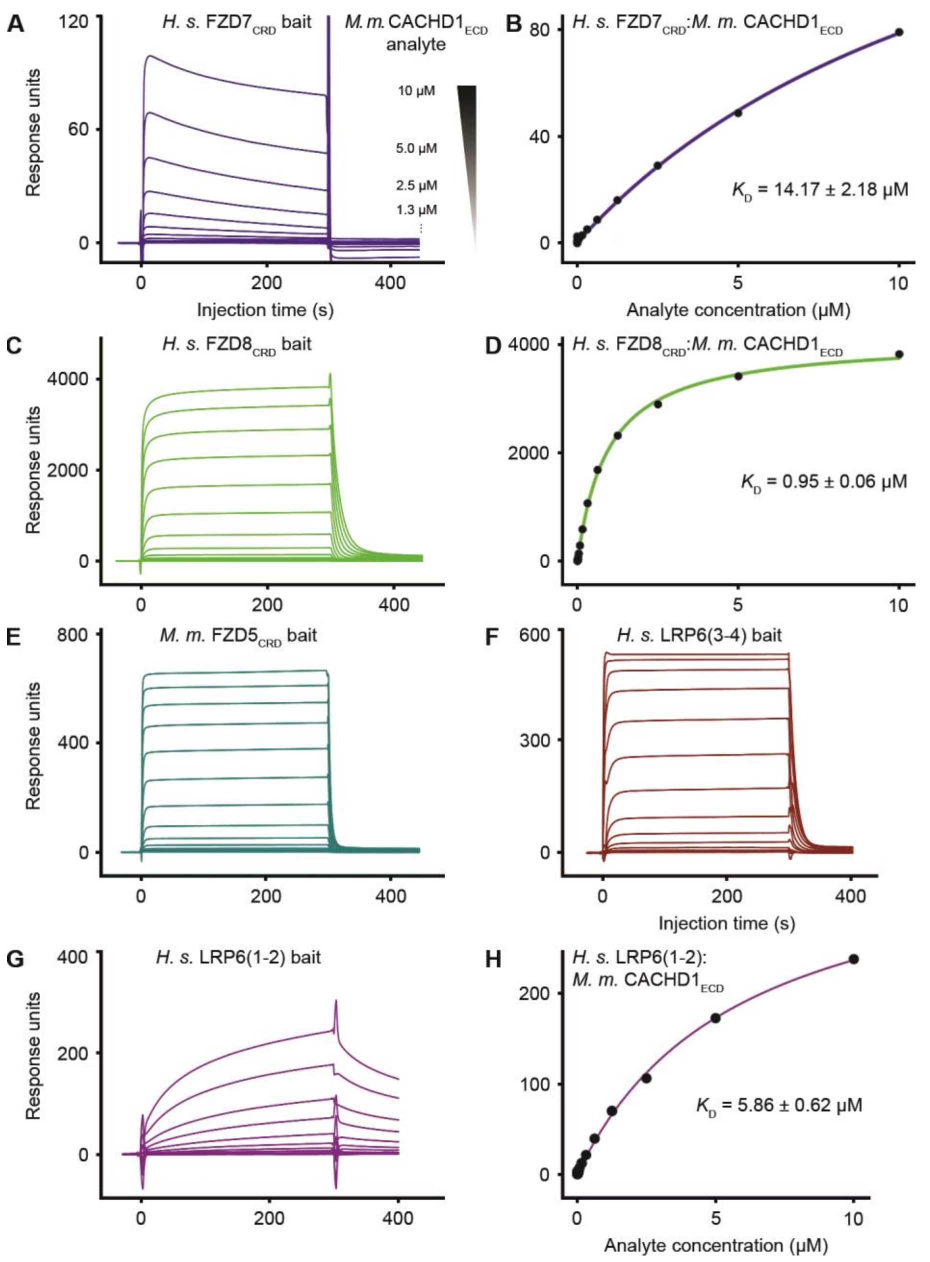


**Fig. S13. SPR analysis of $CACHD1_{ECD}$ interactions with LRP6 and $FZD_{CRD}$ domains.**

(A, C, E-G) Surface plasmon resonance sensorgrams showing the response of different concentrations of mouse $CACHD1_{ECD}$ analyte flowing over surfaces of immobilised human $FZD7_{CRD}$ (A), human $FZD8_{CRD}$ (C), mouse $FZD5_{CRD}$ (E), human $LRP6_{P3E3P4E4}$ (3-4, F), and $LRP6_{P1E1P2E2}$ (1-2, G). (B, D and H) Graphs showing the determination of the equilibrium constants ($K_D$ ± 95% C.I.) for those interactions, except for $FZD5_{CRD}$ and $LRP6_{P3E3P4E4}$ which are shown in Figure 4.

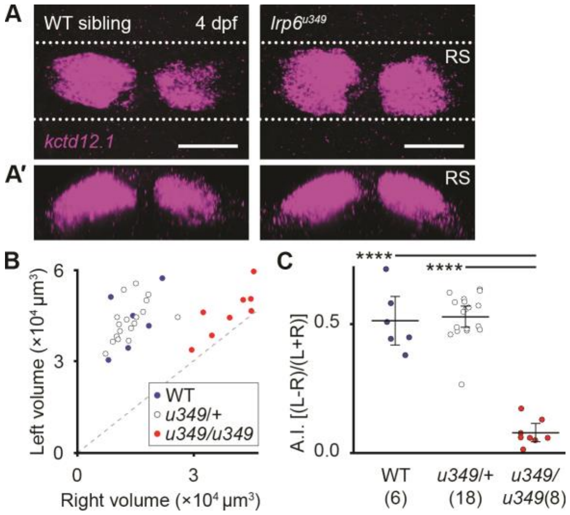


### **Fig. S14. Quantification of *lrp6* mutant phenotype.**

(A) Dorsal and (A′) transverse projection of 4 dpf wildtype sibling and *lrp6*$^{u349}$ mutant larvae after fluorescent RNA *in situ* hybridisation with riboprobes against *kctd12.1*. Maximum projections of confocal stacks. Dotted lines represent the approximate volume in the transverse projections (RS). Scale bars = 50 µm.

(B) Scatterplot showing quantification of *kctd12.1*-fluorescent volumes in the left and right habenulae of wildtype (blue), *u349*/+ (white) and *u349*/*u349* (red) siblings at 4 dpf, represented with a single point. The grey dashed line represents the line of symmetry between left and right volumes. Note the significant increase in the volume of *kctd12.1* in the right habenula of *lrp6*$^{u349}$ mutants compared to wildtype or heterozygous siblings.

(C) Dotplot showing the asymmetry index calculated using *kctd12.1* volumes for each wildtype, *u349*/+ and *u349*/*u349* sibling larvae. Mean asymmetry index for each genotype is indicated with a horizontal bar. Error bars represent 95% confidence intervals of the mean. Wildtype and *lrp6*$^{u349}$ heterozygous siblings have leftward asymmetry of the *kctd12.1* marker, but the increase in right habenula volume renders *lrp6*$^{u349}$ mutants symmetric. ANOVA (degrees of freedom = 2, F = 77.34, $p = 2.38 \times 10^{-12}$) and *post hoc* Tukey pairwise comparisons was used for hypothesis testing, **** $p < 0.005$.

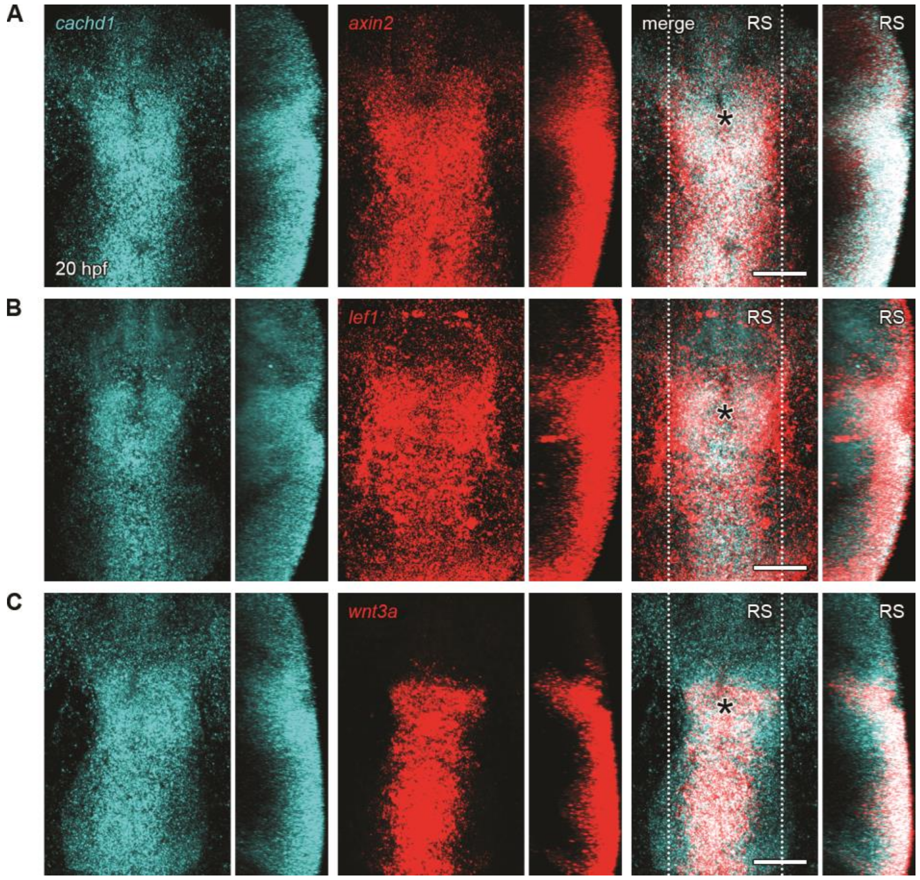


**Fig. S15. *cachd1* is co-expressed with other Wnt pathway genes in the dorsal diencephalon and midbrain roof plate.**

(A-C) Dorsal views (left panels) and sagittal projections (RS, right panels) of ~20 hpf wildtype embryos after double fluorescent *in situ* hybridisation staining with antisense riboprobes for *cachd1* (cyan) and Wnt pathway genes *axin2* (A, red), *lef1* (B, red) or *wnt3a* (C, red) showing *cachd1* is expressed in Wnt active tissues in early development. The approximate position of the pineal is marked with an asterisk. Maximum projections of confocal stacks. Dotted lines represent the approximate volume shown in the sagittal projections (RS).

Scale bars = 50 µm.

### Supplementary tables

#### Table S1. Cross validation of anti-Cachd1 antibody and *cachd1* null allele.

| Genotype | IHC Staining Intensity | | | Total |
|---|---|---|---|---|
| | High | Low | Absent | |
| wildtype | 11 | 2 | 0 | 13 |
| *sa17010*/+ | 0 | 19 | 0 | 19 |
| *sa17010*/*sa17010* | 0 | 1 | 10 | 11 |
| n. d. | 0 | 1 | 0 | 1 |
| Total | 11 | 23 | 10 | 44 |

$\chi^2$ association: $\chi^2 = 70.4$, d. o. f. = 4, $p < 0.001$

**Table S2. Data collection and refinement statistics for the Cachd1:FZD5:LRP6 complex.**

| Data collection | | Refinement | |
|---|---|---|---|
| Source | Diamond I03 | Resolution (Å) | 72.33-4.72 |
| Wavelength(Å) | 0.9762 | No. unique reflections | 23489(154) |
| Space group | $C2_1$ | $R_{work}$ / $R_{free}$ | 0.196/0.243 |
| Cell dimensions: | | No. atoms: | 41757 |
| a, b, c (Å) | 283.70, 198.24, 218.82 | Protein | 41211 |
| α, β, γ (°) | 90, 128.08, 90 | Ligand/ion | 546 |
| Resolution (Å) | 172.26-4.72(4.87) * | Water | 0 |
| $R_{sym}$ or $R_{merge}$ | 0.24(---) | *B*-factors: | |
| $I / \sigma I$ | 6.3(1.6) | Protein | 211.42 |
| Completeness (%) | 88.8(68.0) | Ligand/ion | 216.30 |
| Redundancy | 6.8(7.0) | Water | n/a |
| CC(1/2) | 0.99(0.44) | R.m.s. deviations: | |
| | | Bond lengths (Å) | 0.004 |
| | | Bond angles (°) | 0.86 |

* Values in parentheses are for highest-resolution shell.

**Table S3. Allelic series of *lrp6* nonsense mutations show a bilateral 'double left' phenotype.**

| *lrp6* allele number | Genomic lesion | Freq. of double left phenotype* | | | | $\chi^2$ (2 d. o. f.) | *p* value |
|---|---|---|---|---|---|---|---|
| | | WT | +/- | -/- | Total | | |
| *u348* | 11 bp del. | 0/27 | 2/44 | 27/29 | 100 | 81.58 | $1.93 \times 10^{-18}$ |
| *u349* | 4 bp ins. | 0/17 | 0/55 | 18/19 | 91 | 69.28 | $9.02 \times 10^{-16}$ |
| *u350* | 23 bp del. | 0/25 | 0/46 | 26/27 | 98 | 88.32 | $6.64 \times 10^{-20}$ |
| *u351* | 149 bp del.+9 bp ins. | 0/12 | 0/16 | 9/9 | 37 | 33.52 | $5.27 \times 10^{-8}$ |

* assessed by colorimetric *in situ* hybridisation with *kctd12.1* riboprobe

**Table S4. Oligonucleotides for generating CRISPR/Cas9 sgRNAs targeting *lrp6* exon 2.**

| Gene | Guide | Primer sequence (target sequence in blue) |
|---|---|---|
| *lrp6* | sg1 | GCGTAATACGACTCACTATA**GGCCAACGCCACGCTGGTGA**GTTTTAGAGCTAGAAATAGCAAG |
| *lrp6* | sg2 | GCGTAATACGACTCACTATA**GGCCAGACCGGAGATGACGG**GTTTTAGAGCTAGAAATAGCAAG |

Template oligo sequence: AAAAGCACCGACTCGGTGCCACTTTTTCAAGTTGATAACGGACTAGCCTTATTT TAACTTGCTATTTCTAGCTCTAAAAC

**Table S5. Primer sequences used for mapping, genotyping, qPCR, headloop PCR and cloning.**

| Species | Gene | Name | Sequence |
|---|---|---|---|
| Zebrafish | *ak4* | e1-Mapping-F | CTGTTTTGCACCTCCAACCT |
| Zebrafish | *ak4* | e1-Mapping-R | GCTTCACGGAGCATATGACA |
| Zebrafish | *cachd1* | i8-9-Mapping-F | TTTCAACACTTTGGCCTGTT |
| Zebrafish | *cachd1* | i8-9-Mapping-R | GCAGTGCAGAAGAGGGTTTC |
| Zebrafish | *cachd1* | u761-AloI-F | TTTAACTGCACTGTTTTGCCTTA |
| Zebrafish | *cachd1* | u761-AloI-R | ATAGGCATAAACGGCGAACA |
| Zebrafish | *cachd1* | HRMA-sa17010-F | GAGCAATCTGGAGCTGGGTT |
| Zebrafish | *cachd1* | HRMA-sa17010-R | TGTATGTCGCGGCAGTAAGG |
| Zebrafish | *cachd1* | cloning-FL-SalI-F | ATGAACGTCGACTGCGAAACGGAAAAGTTAGG |
| Zebrafish | *cachd1* | cloning-FL-SacII-R | TGCAATCCGCGGGCACTCAGCGTCCACACT |
| Zebrafish | *cachd1* | RT-PCR-e1-F* | TGCGAAACGGAAAAGTTAGG |
| Zebrafish | *cachd1* | RT-PCR-e9-R | CCCACTGTGGTCTCCAGATT |
| Zebrafish | *cachd1* | RT-PCR-e8-F | CGCAGTGAAAGAGGAGAACC |
| Zebrafish | *cachd1* | RT-PCR-e17-R | CCGAATTTTGTTGCCTTTGT |
| Zebrafish | *cachd1* | RT-PCR-e16-F | CTGGACCGTACCTGGATGTT |
| Zebrafish | *cachd1* | RT-PCR-e27-R* | TGAGGGTTGGTTCTGAGGTC |
| Zebrafish | *lrp6* | CRISPR-sg1-HRMA-F | TGTGTTCCACTGGAGTGACATT |
| Zebrafish | *lrp6* | CRISPR-sg1-HRMA-R | CAGGCCCTGCGCATAGATATAA |
| Zebrafish | *lrp6* | CRISPR-sg2-HRMA-F | GATCTACTGGAGCGACGTGAG |
| Zebrafish | *lrp6* | CRISPR-sg2-HRMA-R | TCGGAGTCGGTCCAGTAAAGTT |
| Zebrafish | *lrp6* | HLPCR-control-F | AGATGTTTTGAAGAGTGCGGTG |
| Zebrafish | *lrp6* | HLPCR-control-R | TAAACATCCCGAAAACAAGCTGC |
| Zebrafish | *lrp6* | HLPCR-HL-sg1-F | CCATCACCAGCGTGGCGTTGAGATGTTTTGAAGAGTG CGGTG |
| Zebrafish | *lrp6* | HLPCR-HL-sg2-R | CACCGTCATCTCCGGTCTGGTAAACATCCCGAAAACA AGCTGC |
| Zebrafish | *slc18a3b* | riboprobe-cloning-F | GGAGAGCTCGTGCGTAATTC |
| Zebrafish | *slc18a3b* | riboprobe-cloning-R | CACTTAGAGGCGTCCATCGT |
| Zebrafish | *zgc:101731* | riboprobe-cloning-F | GGTGTTGAGCGAGAGTTGGT |
| Zebrafish | *zgc:101731* | riboprobe-cloning-R | TGTTTTCAAACCTTTGACTGG |
| Zebrafish | *aoc1* | riboprobe-cloning-F | ACAACGGGCAGTATTTCGAC |
| Zebrafish | *aoc1* | riboprobe-cloning-R | TTCTGTAGCGCACAGGTTTG |

| | | | |
|---|---|---|---|
| Zebrafish | *kiss1* | riboprobe-transcription-F | ATGCTGCTTACTGTCATATTGATG |
| Zebrafish | *kiss1* | riboprobe-T3-transcription-R | GGATCCATTAACCCTCACTAAAGGGACACCTAAAACATGAAGGCAAATACC |
| Zebrafish | *fzd1* | fusion-PCR-5'-F | TGGCCGACTGGAGACTCTTTC |
| Zebrafish | *fzd1* | fusion-PCR-5'-R | CACTGGAAGCGTCCGCGCTG |
| Zebrafish | *fzd1* | fusion-PCR-3'-F | CTCCCAGCGCGGACGCTTCC |
| Zebrafish | *fzd1* | fusion-PCR-3'-R | CGAAAGCAGAGCTTCACACTGTGG |
| Zebrafish | *fzd1* | cloning-NotI-F | GCGGCCGCCACCATGGCAGCTCGCGCTCTCTTC |
| Zebrafish | *fzd1* | cloning-AscI-R | GACTATGGCGCGCCCACTGTGGTTTCTCCCTGTTTGCTGTTCGCC |
| Zebrafish | *fzd2* | cloning-NotI-F | GCGGCCGCCACCATGGCAGCGAGTGGAAGTGTG |
| Zebrafish | *fzd2* | cloning-AscI-R | GAGTGAGGCGCGCCAACAGTGGTTTCTCCTTGTCC |
| Zebrafish | *fzd3b* | cloning-NotI-F | GCGGCCGCCACCATGGGCTGTGTTGTGGATTTACC |
| Zebrafish | *fzd3b* | cloning-AscI-R | TAGTATGGCGCGCCCGCACTGGTCCCGTTCTCCGG |
| Zebrafish | *fzd4* | cloning-NotI-F | GCGGCCGCCACCATGGCTCGGTTTGAGTTCGGG |
| Zebrafish | *fzd4* | cloning-AscI-R | TAGTTAGGCGCGCCCACAACCGTCTCGTTTCCTTTGCCCGG |
| Zebrafish | *fzd5* | cloning-NotI-F | GCGGCCGCCACCATGGGGAAACCTGCAGACGAG |
| Zebrafish | *fzd5* | cloning-AscI-R | GACGCTGGCGCGCCGACATGTGATGAGGGTGCTGATTTGTG |
| Zebrafish | *fzd7a* | cloning-NotI-F | GCGGCCGCCACCATGGCTTTCCTCAAGATGCAAC |
| Zebrafish | *fzd7a* | cloning-AscI-R | TAGTATGGCGCGCCTACCGTCGTCTCGCCCTGGT |
| Zebrafish | *fzd7b* | cloning-NotI-F | GCGGCCGCCACCATGGCGGTACGGGAAGTTGG |
| Zebrafish | *fzd7b* | cloning-AscI-R | GAGTGAGGCGCGCCCACCGTTGTTTCCCCTTGGTTG |
| Zebrafish | *fzd8a* | cloning-NotI-F | GCGGCCGCCACCATGGAGTGCTACCTGTTGGG |
| Zebrafish | *fzd8a* | cloning-AscI-R | TACTCTGGCGCGCCGACTTGGGACAAAGGCATCTGCTTGGG |
| Zebrafish | *fzd8b* | fusion-PCR-5'-F | CCAGAGCACATGCCAGCGCATCC |
| Zebrafish | *fzd8b* | fusion-PCR-5'-R | GTGCACAGGGCTCGCCAGGAATC |
| Zebrafish | *fzd8b* | fusion-PCR-3'-F | GATTCCTGGCGAGCCCTGTGCAC |
| Zebrafish | *fzd8b* | fusion-PCR-3'-R | TCATCACACACGAGAAAGTGGCATTTGTTTTGGAGG |
| Zebrafish | *fzd8b* | cloning-NotI-F | GCGGCCGCCACCATGGACTCGCCTACACAGGG |
| Zebrafish | *fzd8b* | cloning-AscI-R | GACGCTGGCGCGCCCACACGAGAAAGTGGCATTTGTTTTGG |
| Zebrafish | *fzd9a* | cloning-NotI-F | GCGGCCGCCACCATGGGACATTGCATGAAGATTGGG |
| Zebrafish | *fzd9a* | cloning-AscI-R | TGATCGGGCGCGCCAACATGTGTGGGACTGTCTGTATAG |
| Zebrafish | *fzd9b* | cloning-NotI-F | GCGGCCGCCACCATGGGAAGCTCACCTCTGCAAATTG |
| Zebrafish | *fzd9b* | cloning-AscI-R | TAGCTCGGCGCGCCTACATGTGTGGGACAGTCTGAGTAGG |
| Zebrafish | *fzd10* | cloning-NotI-F | GCGGCCGCCACCATGGTTGCTGCCGGTGTCGG |

| | | | |
|---|---|---|---|
| Zebrafish | *fzd10* | cloning-AscI-R | GAGCGTGGCGCGCCTACACAAGTTGCAGGAGGACCTGCTG |
| Zebrafish | *smo* | cloning-NotI-F | GCGGCCGCCACCATGTCCTCCAAGCGCCCCTGCTCCATT |
| Zebrafish | *smo* | cloning-AscI-R | TGCGCAGGCGCGCCAAAATCTGAGTCAGCATCCAATAGCTCAGC |
| Zebrafish | *gng8* | cloning-PCR-F | CATCATACTAGTGGGCTATAAAACAAAATG |
| Zebrafish | *gng8* | cloning-PCR-R | CATCATGATATCTTCGTTTGTAGAGACCAA |
| Mouse | *Cachd1* | qPCR-F | AGTTCAGCAGCTAGCCAAAAA |
| Mouse | *Cachd1* | qPCR-R | CCATCAAACTCCATCATGGA |
| Mouse | *Ccnd1* | qPCR-F | GCCATCCAAACTGAGGAAAA |
| Mouse | *Ccnd1* | qPCR-R | GATCCTGGGAGTCATCGGTA |
| Mouse | *Axin2* | qPCR-F | TCCAGAGAGAGATGCATCGC |
| Mouse | *Axin2* | qPCR-R | AGCCGCTCCTCCAGACTATG |
| Mouse | *Hprt1* | qPCR-F | TCATGAAGGAGATGGGAGGC |
| Mouse | *Hprt1* | qPCR-R | CCACCAATAACTTTTATGTCCCC |
| Human | *CACHD1* | qPCR-F | CTTAAATTCAGTTCTTGCAG |
| Human | *CACHD1* | qPCR-R | CGTAGATGGGTCTACTGCGG |
| Human | *CCND1* | qPCR-F | CTCCGCCTCTGGCATTTTGG |
| Human | *CCND1* | qPCR-R | TCTCCTTGCAGCTGCTTAG |
| Human | *AXIN2* | qPCR-F | AGTGTGAGGTCCACGGAAAC |
| Human | *AXIN2* | qPCR-R | CTTCACACTGCGATGCATTT |
| Human | *ACTB* | qPCR-F | TTCTACAATGAGCTGCGTGTG |
| Human | *ACTB* | qPCR-R | GGGGTGTTGAAGGTCTCAAA |
| Human | *FZD7* | cloning-NotI-F | GCGGCCGCCACCATGCGAGACCCAGGTGCAG |
| Human | *FZD7* | cloning-AscI-R | GAGTGAGGCGCGCCTACCGCAGTCTCCCCCTTGC |
| Jellyfish | *eGFP* | cloning-AscI-F | TAGTATGGCGCGCCGGGTAGCAAGGGCGAGGAGC |
| Jellyfish | *eGFP* | cloning-BamHI-R | GAGGCAGGATCCTCACTTGTACAGCTCGTCCATGCCG |

*also used for riboprobe cloning of *cachd1*

**Table S6. Source of plasmids used as templates for *smo*- and *fzd-eGFP* flow cytometry, SPR and crystallography constructs.**

| Species | Gene | Construct ID | Source | Vector |
|---|---|---|---|---|
| Zebrafish | *fzd1* | IMAGE 9038402 | Source Biosciences | pCR4-TOPO |
| Zebrafish | *fzd2* | | Prof. Masa Tada (gift) (*64*) | |
| Zebrafish | *fzd3b* | IMAGE 7040422 | Source Biosciences | pExpress-1 |
| Zebrafish | *fzd4* | Synthesised clone ODa20912: XM_005173425 | GenScript | pcDNA3.1+-DYK |
| Zebrafish | *fzd5* | IMAGE 9037464 | Source Biosciences | pCR4-TOPO |
| Zebrafish | *fzd6* | IMAGE 6971142 | Source Biosciences | pCMV-SPORT6.1 |
| Zebrafish | *fzd7a* | | Prof. Masa Tada (gift) (*65*) | |
| Zebrafish | *fzd7b* | IMAGE 5777452 | Source Biosciences | pME18S-FL3 |
| Zebrafish | *fzd8a* | IMAGE 7002555 | Source Biosciences | pExpress-1 |
| Zebrafish | *fzd8b* | IMAGE 6802128 | Source Biosciences | pCMV-SPORT6.1 |
| Zebrafish | *fzd9a* | Synthesised clone ODa11014: XM_003198686 | GenScript | pcDNA3.1+-DYK |
| Zebrafish | *fzd9b* | IMAGE 9038534 | Source Biosciences | pCR4-TOPO |
| Zebrafish | *fzd10* | IMAGE 7042011 | Source Biosciences | pExpress-1 |
| Zebrafish | *smo* | | (*66*) | pCS2+ |
| Mouse | $Cachd1_{ECD}$ | IMAGE 6834428 | Source Biosciences | pYX-Asc |
| Mouse | $Fzd5_{CRD}$ | Synthesised clone | GenScript | pNeo_sec |
| Human | *FZD7* | IMAGE 4549389 | Source Biosciences | pOTB7 |
| Human | $FZD7_{CRD}$ | Synthesised clone | GenScript | pNeo_sec |
| Human | $FZD8_{CRD}$ | Synthesised clone | GenScript | pNeo_sec |
| Human | $LRP6_{P1E1P2E2}$ | IMAGE 40125687 | Source Biosciences | pHL_sec |
| Human | $LRP6_{P3E3P4E4}$ | IMAGE 40125687 | Source Biosciences | pHL_sec |

**Table S7. Riboprobe templates.**

| Gene | Vector | Resistance | Linearization | RNA Polymerase | Reference |
|---|---|---|---|---|---|
| *aoc1* | pCRII-TOPO | Amp, Kan | SpeI | T7 | constructed |
| *slc18a3b* | pCRII-TOPO | Amp, Kan | XhoI | SP6 | constructed |
| *kiss1* | | | | T3 | PCR-amplified |
| *cachd1* | pCRII-TOPO | Amp, Kan | SpeI | T7 | constructed |
| *zgc:101731* | pCR-Blunt II-Topo | Kan, Zeocin | NotI | SP6 | constructed |
| *axin2* | pSport 1 | Amp | Asp718 | SP6 | (*67*) |
| *selenop2* | pBS KS+ | Amp | SalI | T7 | (*68*) |
| *otx5* | pBS | Amp | NotI | T7 | (*5*) |
| *kctd12.2* | pBK-CMV | Kan | BamHI | T7 | (*28*) |
| *kctd8* | pCRII-TOPO | Amp, Kan | XhoI | SP6 | (*28*) |
| *kctd12.1* | pBK-CMV | Kan | EcoRI | T7 | (*5*) |
| *prss1* | pCRII-TOPO | Amp, Kan | XhoI | SP6 | (*69*) |
| *spaw* | pGEMT-EASY | Amp | SpeI | T7 | (*70*) |
| *lefty1* | pBS SK+ | Amp | NotI | T7 | (*71*) |
| *aldh1a3* | pGEMT-EASY | Amp | SalI | T7 | (*72*) |
| *dbx1b* | pCRII-TOPO | Amp, Kan | BamHI | T7 | (*23*) |
| *wnt3a* | pBS | Amp | SmaI | T7 | (*73*) |
| *lef1* | pCR-Blunt II-Topo | Kan, Zeocin | SacI | T7 | (*74*) |

**Table S8. HCR probe sets for zebrafish *cachd1* and *lrp6*.**

| Gene | Amplifier | Name | Sequence |
|---|---|---|---|
| *cachd1* | B1 | cachd1_B1_7 | TTCTTGGATTGTTGGCGCAGGGTAA |
| *cachd1* | B1 | cachd1_B1_8 | AACTGCGGTAAACAGACCAGCATTA |
| *cachd1* | B1 | cachd1_B1_9 | GCGTCAAGGCATAGTTTTAGGCACT |
| *cachd1* | B1 | cachd1_B1_10 | TCCGGACTCCGGTAGGGTTACTAAG |
| *cachd1* | B1 | cachd1_B1_14 | CTGTCCGCGACCGACTCGCCATTGT |
| *cachd1* | B1 | cachd1_B1_15 | TCCACGCCGCGAAGGAGAATCGAGG |
| *cachd1* | B1 | cachd1_B1_44 | AGATTTTATCATGTTCGTCGATGGA |
| *cachd1* | B1 | cachd1_B1_45 | GAACAGTATCTGCTATCGTCAACAC |
| *cachd1* | B1 | cachd1_B1_65 | TGAAGGTGGCCACGTCGATCATTCG |
| *cachd1* | B1 | cachd1_B1_66 | CTCCCATCTGGTCCGCATAAGGCAA |
| *cachd1* | B1 | cachd1_B1_82 | CATAAACTATGCAGAGGATAAACGA |
| *cachd1* | B1 | cachd1_B1_83 | GCTGCTTCACCGGGATCTCTGGCTG |
| *cachd1* | B1 | cachd1_B1_86 | GAAGGCAGGAGCTGGGCTGCCCCAG |
| *cachd1* | B1 | cachd1_B1_87 | TCTCGACTGTAGCGAGCTGTTTAAA |
| *cachd1* | B1 | cachd1_B1_88 | TACCAGCGGACAACATCACCGTAGG |
| *cachd1* | B1 | cachd1_B1_89 | TCAGGTGCTCATAGGGAGAGGAGAA |
| *cachd1* | B1 | cachd1_B1_91 | TGTCGCTCAGGTAAGCAGTGTAGTG |
| *cachd1* | B1 | cachd1_B1_92 | GGCCGGGGTTGGCTATAAGTCGAGT |
| *cachd1* | B1 | cachd1_B1_93 | TCACCTCATTCCTCACAGAAGACTT |
| *cachd1* | B1 | cachd1_B1_94 | ATTCATCAGTCACGTGGCTGGTGGC |
| *cachd1* | B1 | cachd1_B1_96 | TGTAACGCCTCACAATGTAGCAGTT |
| *cachd1* | B1 | cachd1_B1_97 | TCCGCAGCACCCCATTGGGCGTTGC |
| *cachd1* | B1 | cachd1_B1_98 | CTTTGTCCATGAGTGAACCGGGGTA |
| *cachd1* | B1 | cachd1_B1_99 | ACCATTGCCTCCTGGTGGGATCGAA |
| *cachd1* | B1 | cachd1_B1_103 | TAGGGGCGTGGATGGTGTGGCTAAT |
| *cachd1* | B1 | cachd1_B1_104 | CTGTGTAACCAGAGGCCATTTGGGA |
| *cachd1* | B1 | cachd1_B1_109 | GGGCCACCAAATAGCCTCTGTCCTC |
| *cachd1* | B1 | cachd1_B1_110 | GACCCTTCGGATCAATCAGTGTCGG |
| *cachd1* | B1 | cachd1_B1_120 | AGAAGGCCAGCGCATCACACGTCTC |
| *cachd1* | B1 | cachd1_B1_121 | AGAGACGGTCCACAGTACTGCAAGC |
| *cachd1* | B1 | cachd1_B1_126 | AAGGCTCCTGGTGCACGTCACAGCT |
| *cachd1* | B1 | cachd1_B1_127 | GACTGGGCTCAATTACAGTCAAAGA |
| *cachd1* | B1 | cachd1_B1_134 | CATAAGGACTCTTGGCGCCCACTAT |

| | | | |
|---|---|---|---|
| *cachd1* | B1 | cachd1_B1_135 | CCTCATCTAAAATGCCCATTCCATC |
| *cachd1* | B1 | cachd1_B1_139 | GGTGCCTATAGGCATAAACTGCCAA |
| *cachd1* | B1 | cachd1_B1_140 | TGTGCTGATGACTGCGACGGTGGAT |
| *cachd1* | B1 | cachd1_B1_150 | CCGCTGAGAGAGGATCGTTATTGCA |
| *cachd1* | B1 | cachd1_B1_151 | CCTCGTCGTGATTGCCCACATCAAC |
| *cachd1* | B1 | cachd1_B1_168 | ACCTTTACGGGCCTGAGAAAAGCTG |
| *cachd1* | B1 | cachd1_B1_169 | ACAAAACCAGGGCTGGGTACTAAAA |
| *lrp6* | B5 | lrp6_B5_9 | TAAACAGCGTCCGTTTAATAGACTC |
| *lrp6* | B5 | lrp6_B5_10 | TCTGAACGCCGCTGGGCGCCGAGCC |
| *lrp6* | B5 | lrp6_B5_20 | CCACGATGACAGAGCGCAGCGACCC |
| *lrp6* | B5 | lrp6_B5_21 | GGCCGTTGGGCCAGTAGATCTCCGT |
| *lrp6* | B5 | lrp6_B5_31 | GCTGGCTGTAGACGTGGATGTCCAT |
| *lrp6* | B5 | lrp6_B5_32 | GGCTCGCCACGTCCATGGGCTGGCG |
| *lrp6* | B5 | lrp6_B5_35 | TTGGACAGGCGCACTGATAGTAGGG |
| *lrp6* | B5 | lrp6_B5_36 | TGTGGTCCTCCAGCAGCTGTACGCC |
| *lrp6* | B5 | lrp6_B5_38 | CTGTGCGGCGCGCTAGCAGGAGGAG |
| *lrp6* | B5 | lrp6_B5_39 | GCGTGTCCAGAGAGATCCGGCGCAG |
| *lrp6* | B5 | lrp6_B5_45 | GCGAGGTCACCACCAGCTGAGCGTC |
| *lrp6* | B5 | lrp6_B5_46 | CGGCGATGCCGTCCGGGTGGTTCAC |
| *lrp6* | B5 | lrp6_B5_66 | TCACCCCGAAGGTCTGGTGGACGAA |
| *lrp6* | B5 | lrp6_B5_67 | AGCCGCCATTAGCCCACGCACACGG |
| *lrp6* | B5 | lrp6_B5_87 | CCATCGCTGCACGGTCTATCTTGGG |
| *lrp6* | B5 | lrp6_B5_88 | GCACCAGAGTGATGCGGCCCGACCC |
| *lrp6* | B5 | lrp6_B5_113 | CGAGGTTTGAGCCGCCAACAGCGCC |
| *lrp6* | B5 | lrp6_B5_114 | CGATGCTCAGGTCGTACGGCTGCAG |
| *lrp6* | B5 | lrp6_B5_126 | CGCTGCTCTCGATGCGCCGCAGGTC |
| *lrp6* | B5 | lrp6_B5_127 | TCACAATCCGATTGGCTCCGGACAG |
| *lrp6* | B5 | lrp6_B5_132 | GCGCCTGTATTTTGGTTCGTCCCTC |
| *lrp6* | B5 | lrp6_B5_133 | CGTGGATGTCGCTCAGTGAGGCGAT |
| *lrp6* | B5 | lrp6_B5_141 | GGATACAGTCCACCTCACCCGACAC |
| *lrp6* | B5 | lrp6_B5_142 | CAAACCCGTCACAGCGCCACGCCTG |
| *lrp6* | B5 | lrp6_B5_147 | CGTCGGAGCGGTCCTGGCAGTTGAT |
| *lrp6* | B5 | lrp6_B5_148 | CAGGGCACAGAACTTCACACTTGTT |
| *lrp6* | B5 | lrp6_B5_152 | CTGTCGCATAGCAGCCGATCTCGTC |
| *lrp6* | B5 | lrp6_B5_153 | TGTTAGTGGGAGCAAACGACGGCTC |
| *lrp6* | B5 | lrp6_B5_155 | ACACCGCGCCGACCACGAACAGCAC |

| | | | |
|---|---|---|---|
| *lrp6* | B5 | lrp6_B5_156 | GGCAGAGCACGCGCTGGCACACGAA |
| *lrp6* | B5 | lrp6_B5_158 | GTCCGTGAACCACGAAGTCATTGGT |
| *lrp6* | B5 | lrp6_B5_159 | GGACGTATCCCAGCGGCACCGGCGG |
| *lrp6* | B5 | lrp6_B5_162 | CTCCCATGATGCTCAGCGAGCCCAT |
| *lrp6* | B5 | lrp6_B5_163 | CGCGGTCGTACGGTGGTCCACTGCT |
| *lrp6* | B5 | lrp6_B5_170 | CGAAGTGGCGGTAACTGTACGGCCG |
| *lrp6* | B5 | lrp6_B5_171 | CCGTGCTGCACGGCGTCGTCGGAGG |
| *lrp6* | B5 | lrp6_B5_175 | GCAGCGGCTCCGAGTCGTAGTTCAG |
| *lrp6* | B5 | lrp6_B5_176 | ACTGGCTGCGCGGCGTGGGCGGCGG |
| *lrp6* | B5 | lrp6_B5_178 | GCTCGGTGTACGGTGACGGCGGGCA |
| *lrp6* | B5 | lrp6_B5_179 | GCGGGTACAGCTGGTGCGAGTAGCT |
